# Single-cell multi-omics reveals dynamics of purifying selection of pathogenic mitochondrial DNA across human immune cells

**DOI:** 10.1101/2022.11.20.517242

**Authors:** Caleb A. Lareau, Sonia M. Dubois, Frank A. Buquicchio, Yu-Hsin Hsieh, Kopal Garg, Pauline Kautz, Lena Nitsch, Samantha D. Praktiknjo, Patrick Maschmeyer, Jeffrey M. Verboon, Jacob C. Gutierrez, Yajie Yin, Evgenij Fiskin, Wendy Luo, Eleni Mimitou, Christoph Muus, Rhea Malhotra, Sumit Parikh, Mark D. Fleming, Lena Oevermann, Johannes Schulte, Cornelia Eckert, Anshul Kundaje, Peter Smibert, Ansuman T. Satpathy, Aviv Regev, Vijay G. Sankaran, Suneet Agarwal, Leif S. Ludwig

**Affiliations:** Department of Pathology, Stanford University, Stanford, CA 94305, USA; Parker Institute of Cancer Immunotherapy, San Francisco, CA 94129, USA; Department of Genetics, Stanford University, Stanford, CA 94305, USA; Broad Institute of MIT and Harvard, Cambridge, MA 02142, USA; Division of Hematology / Oncology, Boston Children’s Hospital and Department of Pediatric Oncology, Dana-Farber Cancer Institute, Harvard Medical School, Boston, MA 02115, USA; Berlin Institute of Health at Charité – Universitätsmedizin Berlin, Charitéplatz 1, 10117 Berlin, Germany; Max-Delbrück-Center for Molecular Medicine in the Helmholtz Association (MDC), Berlin Institute for Medical Systems Biology (BIMSB), 10115 Berlin, Germany; Institute of Biotechnology, Technische Universität Berlin, Berlin, Germany; Department of Biology, Chemistry, Pharmacy, Freie Universität Berlin, Berlin, Germany; Technology Innovation Lab, New York Genome Center, New York, NY 10013, USA; Paulson School of Engineering and Applied Sciences, Harvard University, Cambridge, MA 02134, USA; Center for Pediatric Neurosciences, Mitochondrial Medicine, Cleveland Clinic, Cleveland, OH 44195, USA; Department of Pathology, Boston Children’s Hospital, Harvard Medical School, Boston, MA 02115, USA; Department of Pediatric Oncology, Charité-Universitätsmedizin Berlin, Campus Virchow Klinikum, 13353 Berlin, Germany; Department of Computer Science, Stanford University, Stanford, CA 94305, USA; Department of Biology and Koch Institute, Massachusetts Institute of Technology, Cambridge, MA 02139, USA

## Abstract

Cells experience intrinsic and extrinsic pressures that affect their proclivity to expand and persist *in vivo*. In congenital disorders caused by loss-of-function mutations in mitochondrial DNA (mtDNA), metabolic vulnerabilities may result in cell-type specific phenotypes and depletion of pathogenic alleles, contributing to purifying selection. However, the impact of pathogenic mtDNA mutations on the cellular hematopoietic landscape is not well understood. Here, we establish a multi-omics approach to quantify deletions in mtDNA alongside cell state features in single cells derived from Pearson syndrome patients. We resolve the interdependence between pathogenic mtDNA and lineage, including purifying selection against deletions in effector/memory CD8 T-cell populations and recent thymic emigrants and dynamics in other hematopoietic populations. Our mapping of lineage-specific purifying selection dynamics in primary cells from patients carrying pathogenic heteroplasmy provides a new perspective on recurrent clinical phenotypes in mitochondrial disorders, including cancer and infection, with potential broader relevance to age-related immune dysfunction.

## Introduction

Mitochondria are unique and complex organelles essential for metabolism and also carry their own genome. Characterized by a high mutation rate, cell cycle-independent (relaxed) replication, and variable copy number, mitochondrial DNA (mtDNA) possesses a variety of distinct genetic properties compared to nuclear DNA. In human cells, mitochondrial genomes are present in high copy numbers (100s–1,000s), and mutations in mtDNA may vary in their level of heteroplasmy (proportion of mitochondrial genomes carrying a specific variant) in and across individual cells^1,2^. Notably, mtDNA related disorders affect approximately 1 in 4,300 individuals, many of which present with heterogeneous phenotypes, cell-type specific defects and variable severity that may correlate with heteroplasmy of pathogenic mutations^3^. Furthermore, the age-related accumulation of somatically mutated mtDNA molecules in human cells and tissues may contribute to a variety of complex human diseases, including neurodegenerative, neuromuscular, and metabolic disorders^1,2^ as well as cancer phenotypes^4,5^. While germline single nucleotide variants (SNV) have been more comprehensively studied in human tissues, the effects of a second major class of mutations, large mtDNA deletions and their consequences, have been examined to a lesser extent, particularly in individual cells. Notably, single large-scale mtDNA deletions (SLSMD) have been implicated in a continuum of congenital disorders, including Pearson syndrome (PS), Kearns-Sayre syndrome (KSS), and progressive external ophthalmoplegia (PEO)^6^.

Recently, we and others have demonstrated the utility of single-cell genomics approaches for mtDNA genotyping in combination with readouts of cellular states^7,8^. The droplet-based mitochondrial single-cell assay for transposase accessible chromatin by sequencing (mtscATAC-seq) technique has enabled the scalable, concomitant profiling of accessible chromatin and mtDNA in tens of thousands of single cells^9,10^. The application of these approaches has revealed the high prevalence of somatic mtDNA mutations, many of which may further be stably propagated across cell divisions to facilitate retrospective clonal/lineage tracing studies in *ex vivo* derived human and clinical samples^7–9^. Moreover, they facilitate the study of pathogenic mtDNA variants associated with human disease. In patients with mitochondrial encephalomyopathy lactic acidosis and strokelike episodes (MELAS) caused by the mt3243A>G mutation, we demonstrated a previously unappreciated purifying selection against cells with pathogenic mtDNA in hematopoietic lineages, specifically T cells^10^. While the mechanism of this selection has not been established, these findings suggest that pathogenic mtDNA heteroplasmy is not only intrinsically linked to cellular state and its maintenance, but also cell fate, both of which may evolve over an individual’s lifetime given the dynamic features of heteroplasmy. Moreover, the genomic and compensatory molecular responses to pathogenic mtDNA and associated, possibly cell-type specific metabolic alterations continue to be subject of investigation and have been particularly difficult to resolve in a heteroplasmy-dependent manner in primary patient specimens.

Here, we apply and combine a series of multi-omics single-cell approaches and introduce mgatk-del, a novel computational approach to assess heteroplasmy of mtDNA deletions in single cells, to the study of hematopoiesis in patients with PS. PS is defined by SLSMD and phenotypically characterized by sideroblastic anemia and exocrine pancreas dysfunction^11–13^. By examining primary hematopoietic cells of the peripheral blood and the bone marrow (total n=114,696 new primary cell profiles), we reveal the distribution of pathogenic mtDNA deletions in hematopoietic lineages, its depletion in specific cell types and associated alterations in cellular state as assessed by transcriptional, accessible chromatin and protein expression profiling. More broadly, our findings of the selection dynamics in primary immune cell states provide new perspectives on the higher incidence of malignancies^14^ and complex infections^15^ in patients with primary mitochondrial disorders.

## Results

### Single-cell quantification of pathogenic mtDNA deletions

We have previously demonstrated that mtscATAC-seq yields relatively uniform coverage across the mitochondrial genome and can robustly quantify pathogenic variants in single cells^9,10^. Here, we sought to assess this approach for detecting and quantifying large mtDNA deletions that underlie PS and related pathologies. These large mtDNA deletions have been hypothesized to occur due to strand displacement errors in mtDNA replication between the heavy (O_H_) and light (O_L_) origins of replication (**Fig. 1a**)^16^. To examine these deletions in single-cell data, we conducted mixing experiments by pooling *in vitro* cultured fibroblasts derived from two healthy donors and three patients with PS carrying three distinct mtDNA deletions for mtscATAC-seq (**Fig. 1b**). Following sequencing, cells from each donor were demultiplexed using private SNVs (**Fig. 1b; Methods**). Pseudo-bulk summaries of high-quality cells per donor revealed distinct dips in coverage along the mtDNA genome corresponding to the specific deletions at variable levels of heteroplasmy (**Fig. 1c**).

**Figure 1.**
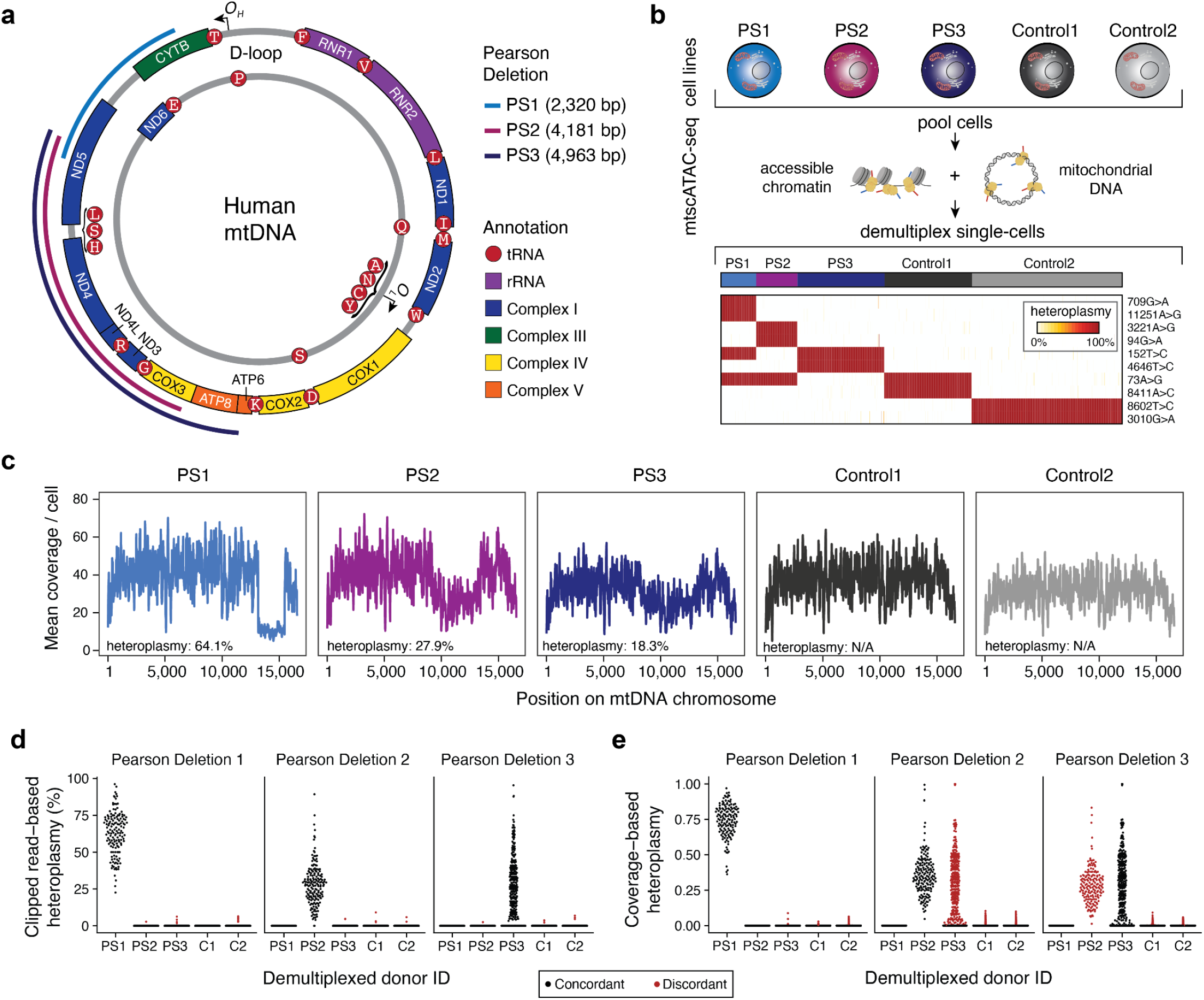
Identification and quantification of heteroplasmic pathogenic mtDNA deletions in single cells. **(a)** Schematic of mitochondrial DNA (mtDNA) in humans with Pearson syndrome (PS) related deletions relevant for the cell lines examined in (b). O_H_ and O_L_ represent the heavy and light chain origins of replication, respectively. PS1, PS2, PS3 represent three different mtDNA deletions identified in three independent donors from which the cell lines were derived. Size and location of deletions as indicated. **(b)** Summary of cell line mixing experiment and demultiplexing using mtDNA haplotype-derived SNVs in single cells. Heatmap depicts homoplasmic SNVs that facilitate the separation of cells from distinct donors. **(c)** Mean coverage plots per cell across the mitochondrial genome for the demultiplexed donor cell identities. Drops in the coverage are indicative of large mtDNA deletions. **(d,e)** Estimates of single-cell heteroplasmy using **(d)** clipped-read enumeration and **(e)** coverage-based approaches. Red dots represent false-positive heteroplasmy assessments from the deletion/donor pair (‘discordant’). Each dot represents a single-cell’s estimated heteroplasmy for each respective deletion. C1, C2 = Control 1 and Control 2 as in (b) and (c).

Though software has been developed to analyze mtDNA deletions in bulk sequencing data^17,18^, these workflows do not ensure valid estimation of deletion heteroplasmy, particularly in lower-coverage libraries such as individual cells, and do not readily scale to thousands of cells from an mtscATAC-seq library. Thus, we developed a computational approach, termed mgatk-del, that utilizes aligned mtDNA sequencing reads that result from the CellRanger-ATAC preprocessing (**Extended Data Fig. 1a; Methods**). To achieve precise heteroplasmy estimation, we reasoned that base-resolution breakpoints in sequencing reads (encoded as soft-clips in the alignments) could be used to infer deletion junctions, which could be corroborated with per-read secondary alignments reported from bwa (**Extended Data Fig. 1b,c; Methods**). Deletion heteroplasmy then could be estimated as a ratio of reads supporting or contradicting a deleted junction sequence. In an effort to benchmark this approach, we observed biases in heteroplasmy estimation as a function of where the deletion junction occurred in the observed sequencing read (**Extended Data Fig. 1c,d**). To account for this, we systematically evaluated values for two key hyper-parameters (‘near-param’ and ‘far-param’; **Extended Data Fig. 1d**) via an exhaustive grid search in a deletion-specific manner after simulating synthetic cells with known levels of heteroplasmy (**Extended Data Fig. 1e; Methods**). Ultimately, this framework enabled us to accurately estimate heteroplasmy for each deletion utilizing clipped reads on chrM from sequencing data.

Having established the computational approach, we quantified single-cell heteroplasmy for all three investigated pathogenic deletions. Following donor demultiplexing, our clipped-read heteroplasmy estimates revealed marked variation in deletion heteroplasmy across the population of cells (**Fig. 1d**), consistent with our prior observations of heterogeneity in cells derived from patients with mitochondrial disorders caused by SNVs ^9,10^. Furthermore, non-zero heteroplasmy was highly specific for each PS cell line (**Fig. 1d**). Conversely, a coverage-based estimate of heteroplasmy (considering ratios of read depths within and outside the deleted region) showed greater non-specific heteroplasmy at deletions discordant from the originating PS patient cells though both methods were overall correlated (**Fig. 1e** and **Extended Data Fig. 1f; Methods**). Together, our analyses demonstrate the development and benchmarking of mgatk-del to enable high-confidence identification of mtDNA deletions and their heteroplasmy in single cells. In contrast to a coverage-based estimate, and in non-mixed settings, mgatk-del heteroplasmy estimates improve the identification of cells with 0% heteroplasmy, thereby uniquely enabling the observation of complete purifying selection of these pathogenic alleles as described below.

### Purifying selection of pathogenic mtDNA deletions in human CD8 T cells

After establishing the use of mtscATAC-seq and mgatk-del to identify and quantify mtDNA deletions, we sought to study these in primary patient cells. We obtained peripheral blood mononuclear cells (PBMCs) from three cases, including a 7-year-old male with PS / Kearns Sayre syndrome (“PT1”), a 4-year-old female with PS (“PT2”), and a 4-year-old male with PS and deletion 7q (del7q) myelodysplastic syndrome (MDS; “PT3”). Each patient presented with a distinct SLSMD. PBMCs from all three patients were profiled using both mtscATAC-seq and 10x 3’ scRNA-seq to quantify heterogeneity of mtDNA deletion heteroplasmy, chromatin accessibility and transcriptional profiles (**Fig. 2a**). Application of mgatk-del revealed the base-resolution breakpoints corresponding to the deleted regions for each patient, without prior knowledge of the deletion breakpoints, and quantification of deletion heteroplasmy in single cells (**Fig. 2b** and **Extended Data Fig. 2a; Methods**). Among cells passing quality control (minimum 1,000 fragments; 45% reads in peaks; 10 reads spanning the deletion junction; total n=15,064), we observed marked variation of heteroplasmy in PBMCs, including hundreds of cells that had no detectable mtDNA deletion heteroplasmy in each of the three patients investigated (**Fig. 2c; Methods**).

**Figure 2.**
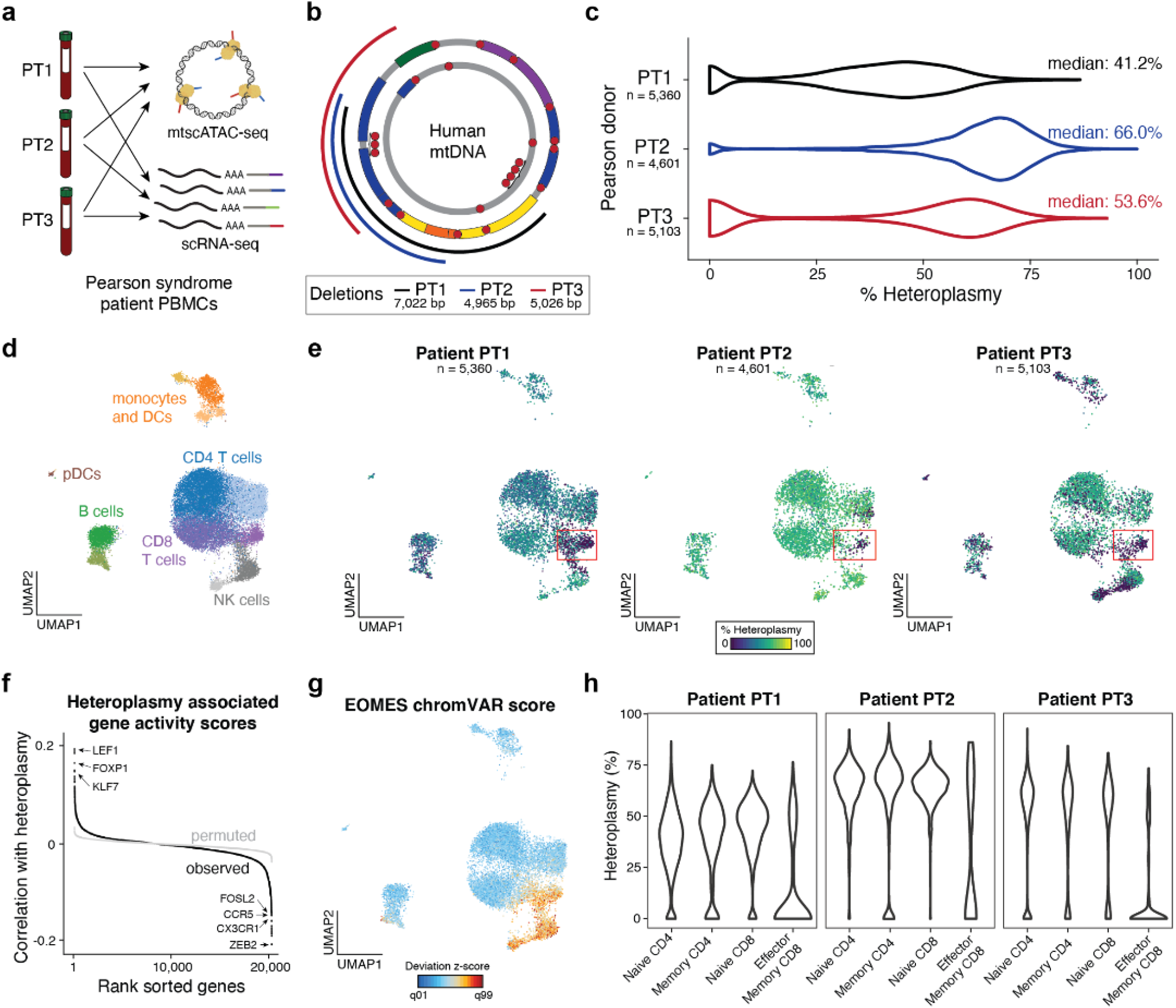
Purifying selection against pathogenic mtDNA deletions in peripheral blood CD8 T cells in Pearson syndrome. **(a)** Schematic of single-cell genomics data generation. Peripheral blood mononuclear cells (PBMCs) from three different Pearson syndrome (PS) patients (PT1, PT2, PT3) were collected and processed via scRNA-seq and mtscATAC-seq. **(b)** Depiction of mtDNA deletions from three investigated PS patients as determined by mgatk-del. Location and size of deletions are indicated. **(c)** Violin plots of single-cell heteroplasmy across indicated patients and respective mtDNA deletions as indicated in (b). Median heteroplasmy (%) and profiled cell numbers are indicated for each patient. **(d)** Reduced dimensionality projection and joint clustering of PBMCs from three PS patients and one healthy control is shown. Major cell types and clusters are annotated in the same color. DC = dendritic cells, pDCs = plasmacytoid dendritic cells, NK = natural killer cells. **(e)** Reduced dimensionality projection as in (d) split by PS patient and colored by respective mtDNA deletion heteroplasmy. The red box indicates a cluster of cells depleted (at 0%) of the respective deletions. **(f)** Correlation of per-gene activity scores in T cells and heteroplasmy across three donors. Top genes are annotated and a permuted background is shown in gray. Genes are shown in rank-sorted order. Select genes have been indicated. **(g)** Annotation of EOMES chromVAR deviation score in single cells. Projection as in (d). **(h)** Violin plots of different CD4+ and CD8+ T cell subsets indicating variable mtDNA deletion heteroplasmy across all three PS patients.

As mtDNA genotypes are paired with single-cell chromatin accessibility data, we sought to examine heteroplasmy variability and dynamics as a function of cell state. We performed dimensionality reduction and clustering using Signac^19^ on PS-derived and healthy donor PBMCs (n=11,325). Our analysis revealed twelve major cell type clusters discernable by gene activity scores, including multiple subpopulations of CD4 and CD8 T cells (**Fig. 2d** and **Extended Data Fig. 2b,c; Methods**). Annotating PS mtDNA deletion heteroplasmy for each of these cells, revealed a cluster of CD8 T cells consistently depleted of heteroplasmy across all three donors (red box; **Fig. 2e**). To better define this population, we performed a systematic correlation analysis of gene activity scores against heteroplasmy within T cells (**Fig. 2f**; **Methods**), revealing that genes most associated with higher heteroplasmy (positive correlation) included *LEF1, FOXP1*, and *KLF7,* genes associated with a naive T cell phenotype ^19,20^. Conversely, genes such as *ZEB2, CX3CR1, CCR5*, and *FOSL2* were strongly associated with lower heteroplasmy and reflective of an effector/memory T cell phenotype^21^. Of note, in specific contexts, *KLF7* has been described to regulate mitochondrial biogenesis^22^ and may be involved in the regulation of oxidative stress^23^.

Further analysis revealed the population of cells with 0% heteroplasmy to have high activity of the EOMES transcription factor (**Fig. 2g**), a factor essential for establishing effector and memory CD8 T cells ^24^. Analysis of heteroplasmy across T cell subsets of all three patients showed consistent depletion of mtDNA deletions in effector/memory CD8 T cells (CD8.TEM) but no other cell state (**Fig. 2h** and **Extended Data Fig. 2d-g**). These results are reminiscent of the previously described purifying selection of pathogenic mtDNA in T cells from patients with MELAS^10^ but adds an additional nuance by revealing a subpopulation of CD8 T cells that depicts this selection in the context of PS. These analyses further underline the utility of mtscATAC-seq^9,10^ and related high-throughput single-cell multi-omic approaches capturing mtDNA^25–27^ in the study of mitochondrial disorders.

### Altered transcriptional states in Pearson syndrome

Next, we leveraged the transcriptomic and chromatin accessibility profiles to discern changes of genomic features associated with PS. To perform these comparisons, we projected the scRNA-seq data from 15,317 PS cells and 19,539 cells from healthy control cells onto a reference PBMC atlas via the Seurat/Azimuth projection (**Fig. 3a**)^28^. This reference-based annotation yielded largely comparable abundances of cell types per donor where the largest difference was ~2x fewer CD14+ monocytes in the PS patients and an overall reduction of CD8^+^ Naive T Cells (**Extended Data Fig. 3a**). Notably, in 3’ scRNA-seq data, we were not able to estimate mtDNA deletion heteroplasmy in individual cells nor in donor-level pseudobulk populations, due to low mitochondrial RNA-derived gene capture and/or potential transcriptional compensation from healthy mtDNA in PS cells (**Extended Data Fig. 3b**). Thus, transcriptional analyses focused on comparing profiles of PS cells to those of healthy controls without further stratification by mtDNA heteroplasmy.

**Figure 3.**
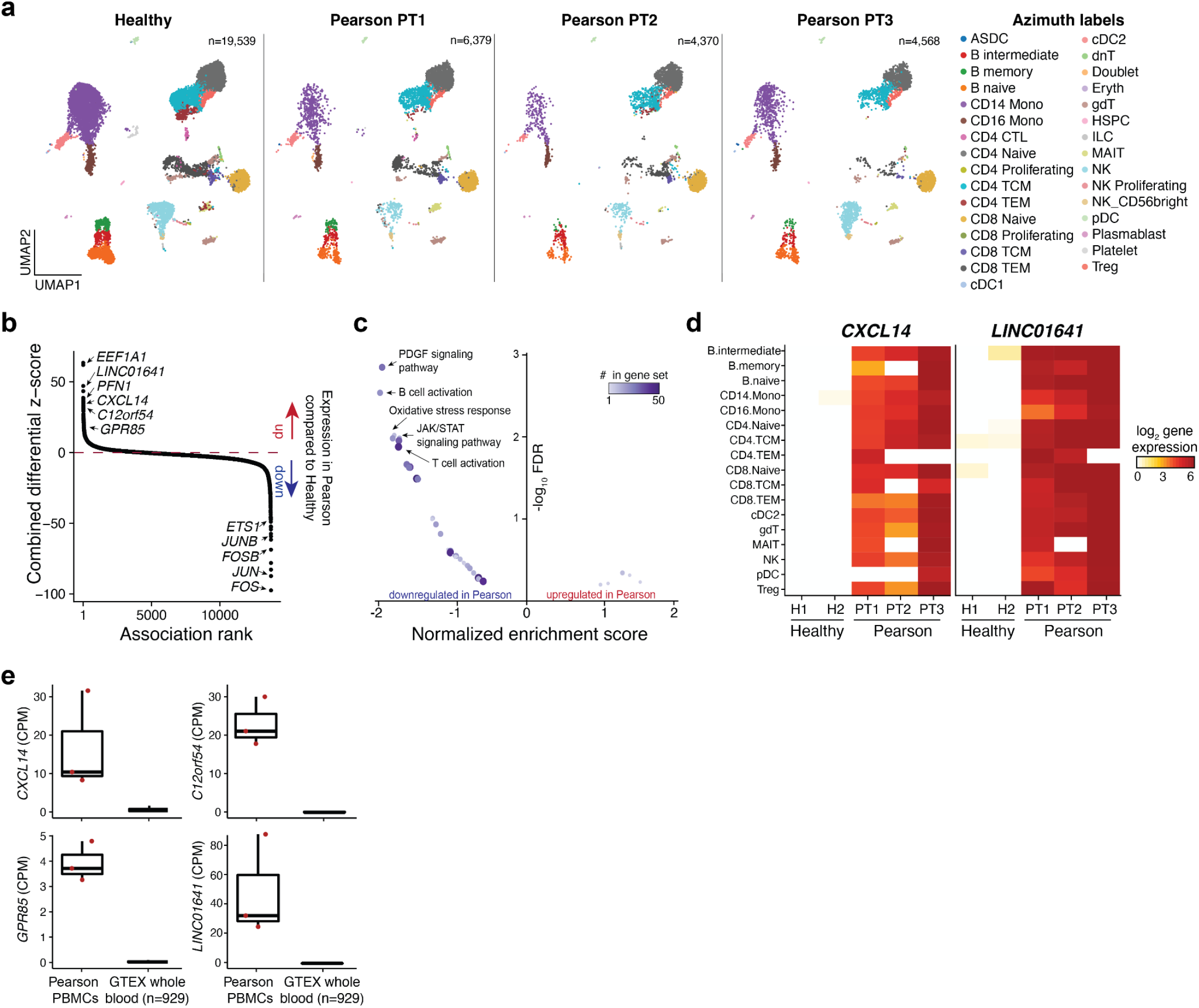
Transcriptional and chromatin accessibility alterations in Pearson syndrome PBMCs. **(a)** Reduced dimensionality projection of three PS patients and healthy controls onto a PBMC reference panel via Azimuth. Full descriptions of abbreviated cell populations are described^28^. **(b)** Rank-ordered combined (across all cell types) differentially expressed genes in PS PBMCs. Selected top genes overexpressed and downregulated in PS are annotated. **(c)** Volcano plot of pathway enrichment analysis results using summary statistics combined across all cell types. **(d)** Heatmap of pseudobulk gene expression across PBMC cell types for indicated genes *CXCL14* and *LINC01641* that are specifically up-regulated in PS compared to healthy controls. **(e)** Expression of genes up-regulated in PS *(CXCL14* and *GPR85)* compared to a larger healthy cohort of whole-blood RNA-seq expression data from GTEx. CPM = counts per million. n indicates the number of donors profiled in GTEx.

For 17 well-represented cell types (>250 cells total between donors), we performed differential gene expression analyses and aggregated summary statistics to identify genes systematically changed in PS compared to controls (**Fig. 3b; Methods**). Though our aggregation prioritized genes with consistent effects across all subtypes of PBMCs, we also observed genes such as *NFAM1* (T cells) and *PLEKHD1* (plasmacytoid dendritic cells; pDCs) that were differentially expressed in only specific cell types (**Extended Data Fig. 3c**). Using our aggregated summary statistic, we performed pathway enrichment analysis that yielded only significantly enriched pathways that were downregulated in PS, including related to PDGF signaling, the oxidative stress response, and JAK/STAT Signaling (**Fig. 3c; Methods**). Many of these pathways, including PDGF signaling as well as B cell and T cell activation, were enriched due to the predominant down-regulation of members of the activator protein 1 (AP-1) complex, including *JUN, FOS, JUNB*, and *FOSB* in PS cells (**Fig. 3b**).

Though we did not observe an enrichment of biological pathways upregulated in PS, we sought to more closely examine up-regulated genes in this condition. In addition to *PFN,* a gene linked to regulating glycolysis and mitochondrial respiration in murine hematopoietic stem cells (HSCs)^29^, we identified *CXCL14, LINC01651,* and *C12orf54* as genes specifically expressed in PS cells (**Fig. 3d** and **Extended Data Fig. 3d**). Analysis of GTEx data indicated that *LINC01651* and *C12orf54* expression is largely restricted to the testis in healthy adults, with otherwise no clearly defined function of these genes in humans (**Extended Data Fig. 3e**), underlining their notable expression in PS PBMCs. Furthermore, we observed increased expression of *GPR85*, the cognate receptor of *CXCL14*^30^, in PS PBMCs, noting that both genes are not normally expressed in blood samples profiled by GTEx (**Fig. 3e** and **Extended Data Fig. 3d,e**) or cord blood^31^. Furthermore, *CXCL14* has been linked to organismal inflammation and pleiotropic functions in glucose metabolism^32^. Overall, these consistently observed PS-specific gene expression signatures may represent candidate biomarkers for PS.

Given the impact of mtDNA deletions on mitochondrial metabolism in PS, we assessed differentially expressed metabolic gene profiles and performed a targeted enrichment analysis of these pathways (**Extended Data Fig. 3f; Methods**). We observed an overall increase of expression of genes encoding mitochondrial-dependent oxidative respiration in most cell states for PS compared to healthy cells. Given the presumed deficiency in oxidative phosphorylation due to the mtDNA deletions, these observations suggest a compensatory transcriptional response in PS. Conversely, we observed decreased expression of genes linked to glycolysis and gluconeogenesis in PS cells. These observations are consistent with recent observations in tumors with loss-of-function mutations in complex I of the respiratory chain^4^ and underline retrograde signaling and gene regulatory mechanisms to compensate for or balance the metabolic alterations resulting from pathogenic mtDNA^33^. Taken together, our analyses point to distinct transcriptional programs in PS, characterized by genes not typically expressed in blood and immune cells and a compensatory metabolic gene expression program.

### Identification of mosaic del7q cells in a patient with Pearson syndrome and Myelodysplastic syndrome

For PT3 with PS, clinical evaluation revealed a mosaic chromosome 7q deletion (del7q), a chromosomal abnormality consistent with the development of myelodysplastic syndrome (MDS) on the backdrop of a congenital bone marrow failure syndrome (**Fig. 4a**)^34^. Notably, the acquisition of monosomy 7 has been recently reported in a case of Pearson syndrome^35^. In the context of PT3, a bone-marrow aspirate enabled the application of mtscATAC-seq to profile bone-marrow mononuclear cells (BMMNCs) with and without CD34+ enrichment. The multi-modal nature of the resulting data thereby readily enabled analysis of the association of mtDNA deletion heteroplasmy and the nuclear del7q abnormality at single-cell resolution. To assess the distribution of del7q cells, we first examined the abundance of fragments overlapping the deleted region, which revealed a clear multimodal distribution in PT3-derived cells, particularly in the CD34+ hematopoietic stem and progenitor cells (HSPCs), compared to healthy control cells (**Fig. 4b** and **Extended Data Fig. 4a**). In addition, we utilized a Gaussian mixture model of gene activity scores as previously implemented^36^, which agreed with our initial inference of del7q status, ultimately annotating cells based on the mixture component probabilities (**Extended Data Fig. 4b,c; Methods**). As the del7q was most abundant in the CD34+ HSPC compartment, we examined the association between del7q and mtDNA heteroplasmy in these cells. Notably, we observed a striking disparity where del7q cells had significantly higher levels of heteroplasmy, suggesting the acquisition of del7q in cells already affected by the pathogenic mtDNA deletion (**Fig. 4c** and **Extended Data Fig. 4d**).

**Figure 4.**
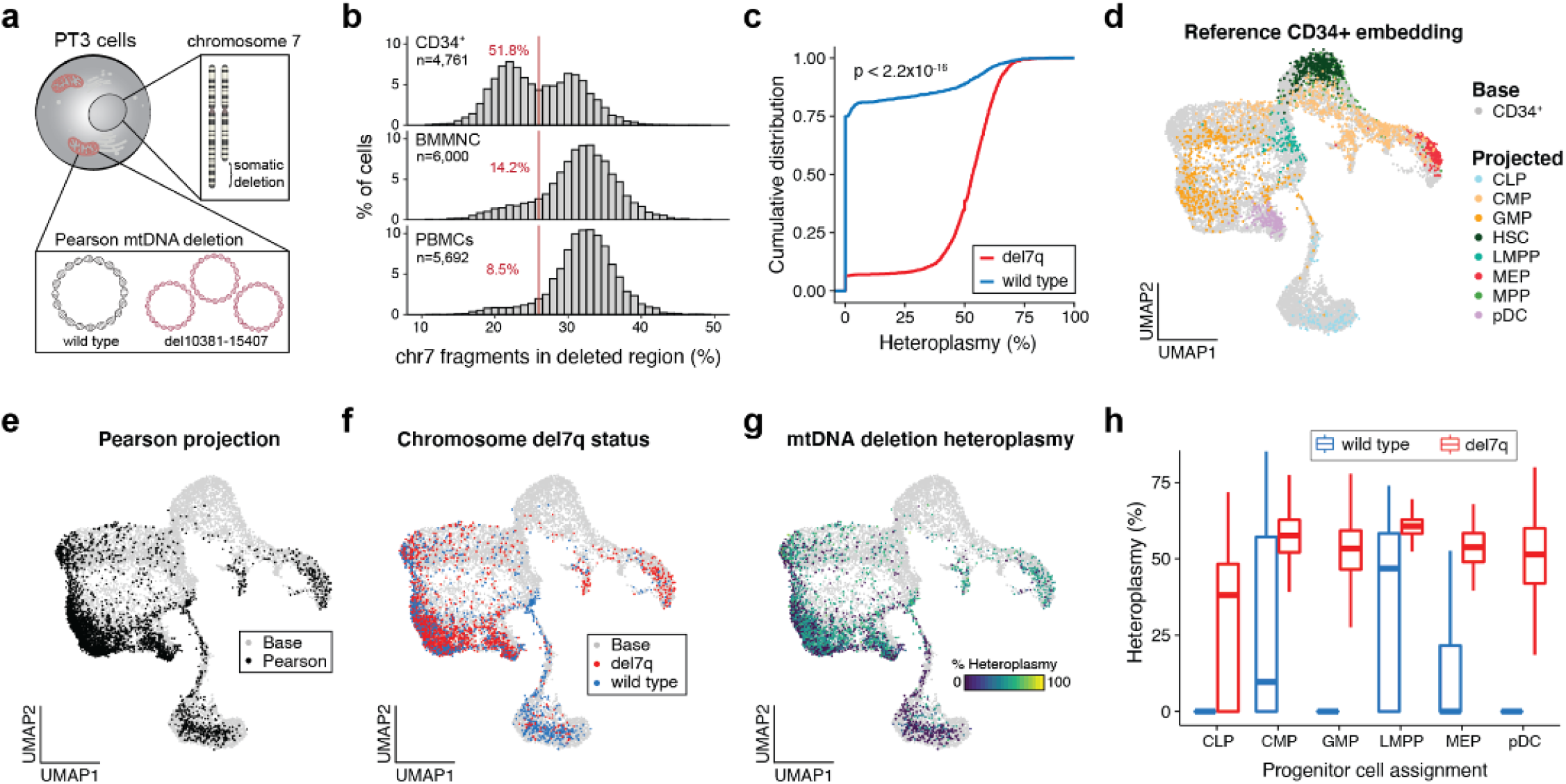
Myelodysplastic cells resolved by a chromosomal 7q deletion in CD34+ hematopoietic stem and progenitor cells in a case of Pearson syndrome. **(a)** Schematic depicting the chromosome 7q deletion (del7q) and mtDNA deletion of PT3 in the same cell. **(b)** Histograms showing the percentage of fragments on chromosome 7 mapping to the somatically deleted region. A consistent cutoff of 26% is shown (red) with the percentage of cells below this threshold being indicated in red. n = cells assayed for each indicated population of CD34+, BMMNC and PBMCs. **(c)** Cumulative distribution curves of mtDNA heteroplasmy stratified by the del7q status per cell. Statistical test: two-sided Kolmogorov-Smirnov test. **(d)** UMAP of a reference CD34+ based embedding (base; grey) with sorted cell populations projected (color-coded) onto the reference. All cells were derived from healthy donors. CLP=common lymphoid progenitor, CMP=common myeloid progenitor, GMP=granulocyte-monocyte progenitors, HSC=hematopoietic stem cell, LMPP=lymphoid primed multipotent progenitor, MEP=megakaryocyte erythroid progenitor, MPP= multipotent progenitor, pDC= plasmacytoid dendritic cell. **(e)** Projection of PS CD34+ cells onto the same base embedding as in (D). **(f)** Annotation of del7q status for each PS CD34+ cell, indicating diploid (blue) and del7q (red) status. **(g)** Annotation of single-cell mtDNA deletion heteroplasmy per PS CD34+ cell. **(h)** Heteroplasmy (%) stratified based on annotated CD34+ progenitor cell state and by del7q ploidy status. Boxplots: center line, median; box limits, first and third quartiles; whiskers, 1.5× interquartile range.

To refine our analysis, we projected CD34+ PT3 data onto a healthy donor reference of sorted CD34+ cells to define the continuous differentiation trajectory of these progenitors via patterns of chromatin accessibility (**Fig. 4d**; **Methods**)^37,38^. Relative to healthy control cells, PT3 displayed a stark depletion of cells annotated as hematopoietic stem cells (HSCs) and multipotent progenitors (MPPs) as well as an enrichment of granulocyte-monocyte progenitors (GMP), resulting in a markedly different estimated composition of the entire HSPC compartment (**Fig. 4e** and **Extended Data Fig. 4e,f**). We observed the pronounced presence of del7q in PT3 GMPs and multipotent erythroid progenitors (MEP), consistent with the MDS phenotype (**Fig. 4f**). Notably, the PT3 common lymphoid progenitor (CLP) population was mostly wild-type for chr7 and depleted of pathogenic mtDNA (**Fig. 4g,h**). Re-analysis of the PBMC data of PT3 (**Fig. 2d,e**) confirmed the presence of del7q carrying cells in peripheral blood, particularly in monocytes **Extended Data Fig. 4g**), whereas del7q-negative cells were mainly located in the lymphocyte compartment, consistent with del7q-positive GMPs and mostly chr7 wild-type CLPs (**Fig. 4f**). Together, these analyses reveal the complexity of lineage commitment and differentiation in the presence of pathogenic mtDNA deletion heteroplasmy and onset of MDS within the early hematopoietic progenitor compartment of this PS patient.

### Multi-omic analysis and purifying selection of pathogenic mtDNA during hematopoietic development

We recently introduced ATAC with Selected Antigen Profiling by sequencing (ASAP-seq), complementing mtscATAC-seq with the concomitant antibody-based quantification of protein expression profiles in single cells ^25^. Applying ASAP-seq to PT3-derived BMMNCs, we profiled 242 surface antigens, yielding 20,580 high-quality cells with quantification across four distinct modalities per cell (chromatin accessibility, nuclear chromosomal aberrations, mtDNA genotypes, surface protein abundance; **Fig. 5a; Methods**). As expected, ASAP-seq retained the capacity to quantify pathogenic mtDNA deletion heteroplasmy in PS cells, revealing variation in heteroplasmy across hematopoietic lineages with the protein measurements facilitating more highly resolved inferences of cell type/state (**Fig. 5b-d** and **Extended Data Fig. 5a; Methods**). Furthermore, our mixture model approach enabled the identification of PT3-derived del7q carrying BMMNCs that were mainly found in the myeloid and erythroid compartments and largely absent in lymphocytes (**Fig. 5e**), consistent with the observed del7q distribution in the CD34+ HSPC and peripheral blood compartments of that same patient (**Fig. 4g**).

**Figure 5.**
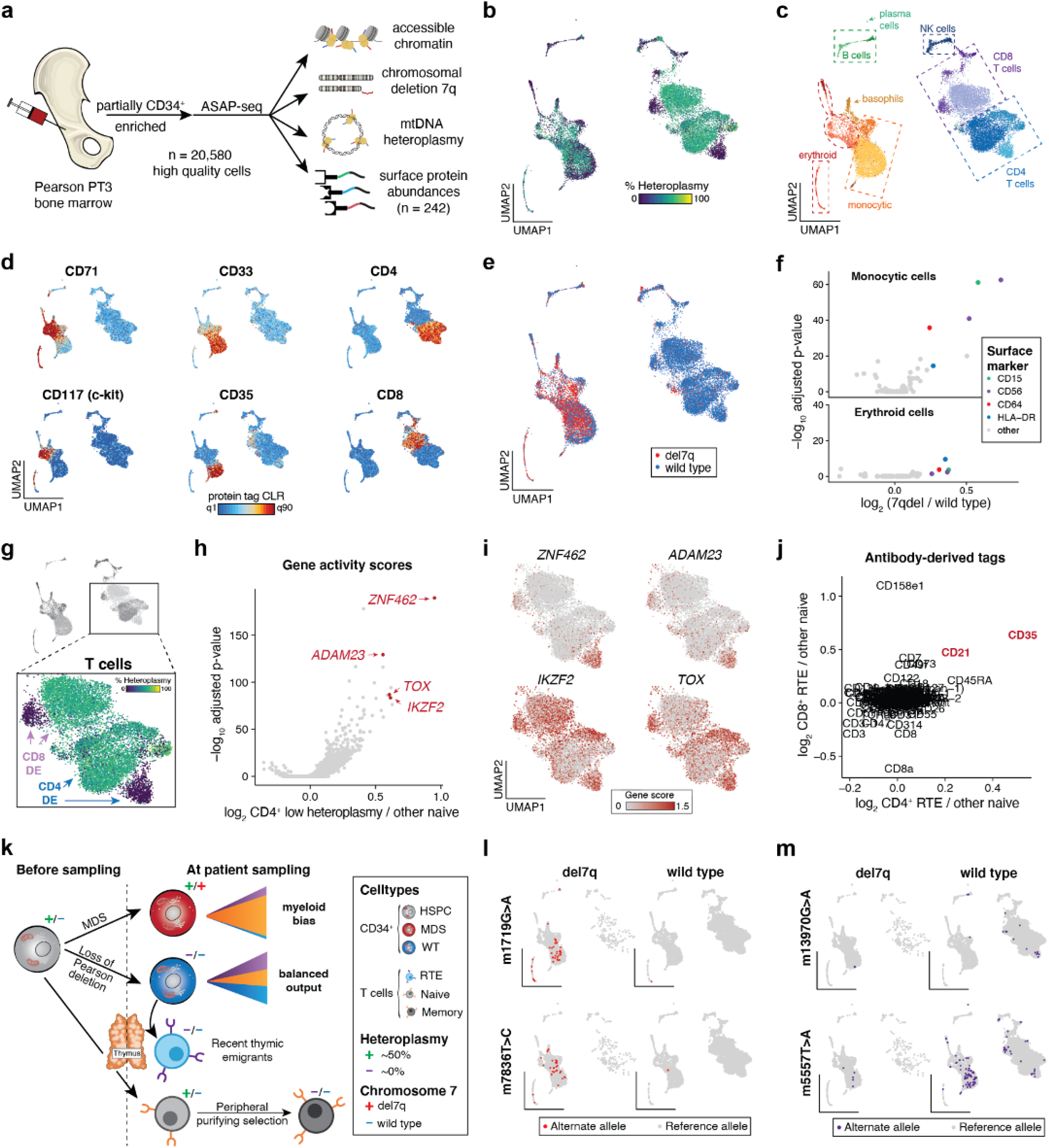
Multi-modal characterization of Pearson syndrome bone marrow mononuclear cells with ASAP-seq. **(a)** Schematic of ASAP-seq experiment from PS bone marrow mononuclear cells (BMMNCs) derived from PT3 with MDS, including the resulting modalities inferred per cell. **(b)** Dimensionality reduction and embedding for high-quality BMMNCs. The corresponding per-cell heteroplasmy is shown across the indicated heteroplasmy color gradient. **(c)** The same embedding as in (b) annotated by major cell type clusters. **(d)** Selected lineage-defining surface protein markers are shown on the reduced dimension space as in (b, c). **(e)** Projection of annotated del7q status onto the UMAP space as in (b, c). **(f)** Volcano plots of differentially expressed protein surface markers inferred from antibody barcodes for del7q versus wild-type cells annotated as erythroid or monocytic from panel (c). **(g)** Schematic illustrating CD4 and CD8 T cell clusters used for differential gene score expression (DE) analyses to identify markers distinguishing low-heteroplasmy cell populations. **(h)** Volcano plot showing differential gene activity scores for comparison of CD4 T cell clusters as illustrated in panel (g). Genes in red *(ZNF462, ADAM23, IKZF2, TOX)* indicate marker genes for recent thymic emigrants (RTEs). (**i**) Projected gene scores for indicated marker genes onto UMAP space as highlighted in (g). **(j)** Differentially expressed proteins for the comparisons of CD4 and CD8 T cell populations as illustrated in panel (g). CD21 and CD35, shown in red, are known surface markers for RTEs. **(k)** Schematic of multi-faceted clonal output and purifying selection in PT3 with PS and MDS. Major cell transitions are depicted as a function of 7qel status and mtDNA deletion heteroplasmy. Notably, selection against cells with high levels of the pathogenic mtDNA deletion appears to occur multiple times: in CD34+ CLPs that do not have the del7q alteration, in RTEs and in the periphery during CD8 T effector/ memory cell formation. **(l)** Projection of somatic mtDNA mutations 1719G>A (15% threshold) and 7836T>C (15%) enriched in cells carrying the del7q. **(m)** Projection of somatic mtDNA mutations 13970G>A (25% threshold) and 5557T>A (35%) enriched in wild-type cells (diploid for chr 7), including in RTEs. For panels (l,m) the threshold for + cell heteroplasmy is based on the empirical density of each variant with the threshold being indicated in parentheses for each variant.

With these distinct modes of measurement, we performed integrative analyses to determine surface markers over-expressed on del7q compared to chr7 wild-type monocytic and erythroid cells (**Fig. 5f; Methods**). For both comparisons, we observed an enrichment of surface proteins CD15, CD56, CD64, and HLA-DR, all markers that have previously been reported to be upregulated in MDS patients^39,40^. Unlike CD56, other markers of NK cells such as CD335 were not expressed on MDS-associated cells, but were present exclusively on NK cells, confirming the specifically altered expression of CD56 in the context of MDS (**Extended Data Fig. 5b**). In addition, we sought to corroborate our observation of the depletion of phenotypic HSCs as revealed in the CD34+ projection analysis (**Fig. 4d,e**). To do this, we compared the distribution of HSPC populations and protein markers in PS to a healthy donor bone marrow specimen previously profiled by ASAP-seq^25^ (**Extended Data Fig. 5c**). Notably, we were unable to detect CD34^+^c-Kit^+^CD71^-^ cells in PT3 despite the clear presence of these cells in healthy BMMNCs (**Extended Data Fig. 5d**). Noting that HSCs do not express CD71, our integrated analysis confirms the apparent relative depletion of phenotypic HSCs in PT3, which may further present a consequence of the pathogenic mtDNA deletion and/or the MDS phenotype.

Next, we investigated two sub-populations of CD4 and CD8 T cells that were notably fully depleted of pathogenic mtDNA heteroplasmy (**Fig. 5g**). To discern their cell state, we performed differential gene score analyses of both populations to other CD4 or CD8 T cells from the same population of BMMNCs, respectively (**Methods**). Our results revealed markers associated with recent thymic emigrants (RTEs)^41^ in both T cell populations, including *ADAM23, IKZF2, TOX,* and *ZNF462* (**Fig. 5g-i** and **Extended Data Fig. 5e**). Differential protein expression analyses of the same populations showed a relative enrichment of CD21 and CD35 in both T cell populations (**Fig. 5k**). Notably, CD21 and CD35 encode the complement receptor (CR) genes CR1 and CR2, which are both upregulated on RTEs^41^ compared to other naive T cells. Notably, we verified the presence of RTE-heteroplasmy depleted cells in peripheral blood by reclustering PBMCs from PT3 (**Extended Data Fig. 5i,j**), which did not separate RTEs in our previous integrated analysis as these were unique to PT3 (**Fig. 2** and **Extended Data Fig. 5k**). In addition, we confirmed our inferences of purifying selection of CD8.TEM cells in the bone marrow, as these cells were similarly depleted for pathogenic mtDNA heteroplasmy and expressed CD195 a marker of T cell activation^42^ (**Extended Data Fig. 5f-h**). Overall, integrating our observations of populations depleted of pathogenic mtDNA deletions, including CLPs in the CD34+ compartment (**Fig. 4d,g**), subpopulations of CD4/CD8 RTE and CD8.TEM cells in the bone marrow and the peripheral blood, ultimately suggest multiple distinct modes of purifying selection of conceivably metabolically impaired cells at distinct stages of lymphopoiesis (**Fig. 5k**).

### mtDNA based clonal tracing in pediatric hematopoiesis and malignancies

As we have previously demonstrated somatic mitochondrial SNVs to identify clonal subsets in hematopoietic populations of adults^7,9^, we sought to determine the prevalence of these mutations at 4 years of age. Application of mgatk with standard parameters revealed 69 somatic mtDNA SNVs that were enriched in expected nucleotide substitution patterns^9,25^ (**Extended Data Fig. 5l**). High allele frequency variants were present in both del7q and wild-type cells (**Extended Data Fig. 5m**), consistent with a model of somatic mtDNA mutations also arising following the acquisition of the nuclear chromosomal abnormality. For example, mutations 1719G>A and 7836T>C refined clones within the del7q compartment (**Fig. 5l**) whereas mutations 12242A>G, 14476G>A, 5557T>A, and 13970G>A were predominantly found in cells with wild type chr7 (**Fig. 5m** and **Extended Data Fig. 5n**). Notably, the 5557T>A mutation was observed in both CD4+ and CD8+ RTEs and in the myeloid compartment, suggesting that the HSPC carrying this mutation is capable of multi-lineage output. Conversely, variants 12242A>G and 14476G>A were only identified in lymphoid cells. Together, our analyses confirm that the application of mtDNA-based lineage tracing extends to pediatric patients and clonal myeloid disorders.

### Altered differentiation but not purifying selection in an in vitro erythroid model

A hallmark feature of PS is severe macrocytic sideroblastic anemia, frequently detectable in the neonatal period, and often in the context of evolution to pancytopenia^11^. Given this clinically significant phenotype, we sought to understand the selection dynamics and altered gene expression programs underlying defective erythropoiesis. To realize this, we *in vitro* differentiated PT3 BMMNCs and healthy control cells in the presence of erythropoietin and other cytokines, collecting cells at day 6 and 12 of culture, before processing with scRNA-seq and mtscATAC-seq (one culture per assay; **Fig. 6a**). To minimize batch effects, PS and healthy cells were pooled for single cell processing and then computationally deconvolved using donor-specific SNPs (**Methods**). Phenotypically, PS cells displayed poor proliferation and clear signs of impairment during erythroid differentiation as assessed by surface markers and cytology (**Extended Data Fig. 6a,b**). Focusing on the PS cells, we estimated per-cell mtDNA deletion heteroplasmy and del7q status to assess the dynamics of these genetic alterations during continued culture and compared the two time points to all BMMNCs (the starting population of the culture) and CD34+ cells (the HSPCs from which most *in vitro* differentiated erythroid cells would most likely arise from). We observed an increase in the proportion of del7q cells already at day 6, indicating that these cells had a proclivity to survive and differentiate *in vitro* (**Fig. 6b**). Furthermore, from day 6 to 12 we did not observe selection against cells carrying the pathogenic mtDNA deletion from either the proportion at 0% (wild type mtDNA) or the overall distribution of cells (**Fig. 6c,d**). Importantly, due to the strong association between del7q and mtDNA deletion heteroplasmy (**Fig. 4c**), we could not conclusively determine the effects of either genetic alteration individually in this experiment. However, our results indicate that the selective pressure during *in vitro* erythropoiesis appears insufficient to yield a similar form of purifying selection of cells with high pathogenic mtDNA heteroplasmy, standing in contrast to our observations of such phenomena in various T cell populations in PS and MELAS^10^. Further studies of *in vivo* erythropoiesis will be required to better understand mtDNA deletion heteroplasmy dynamics during erythroid differentiation.

**Figure 6.**
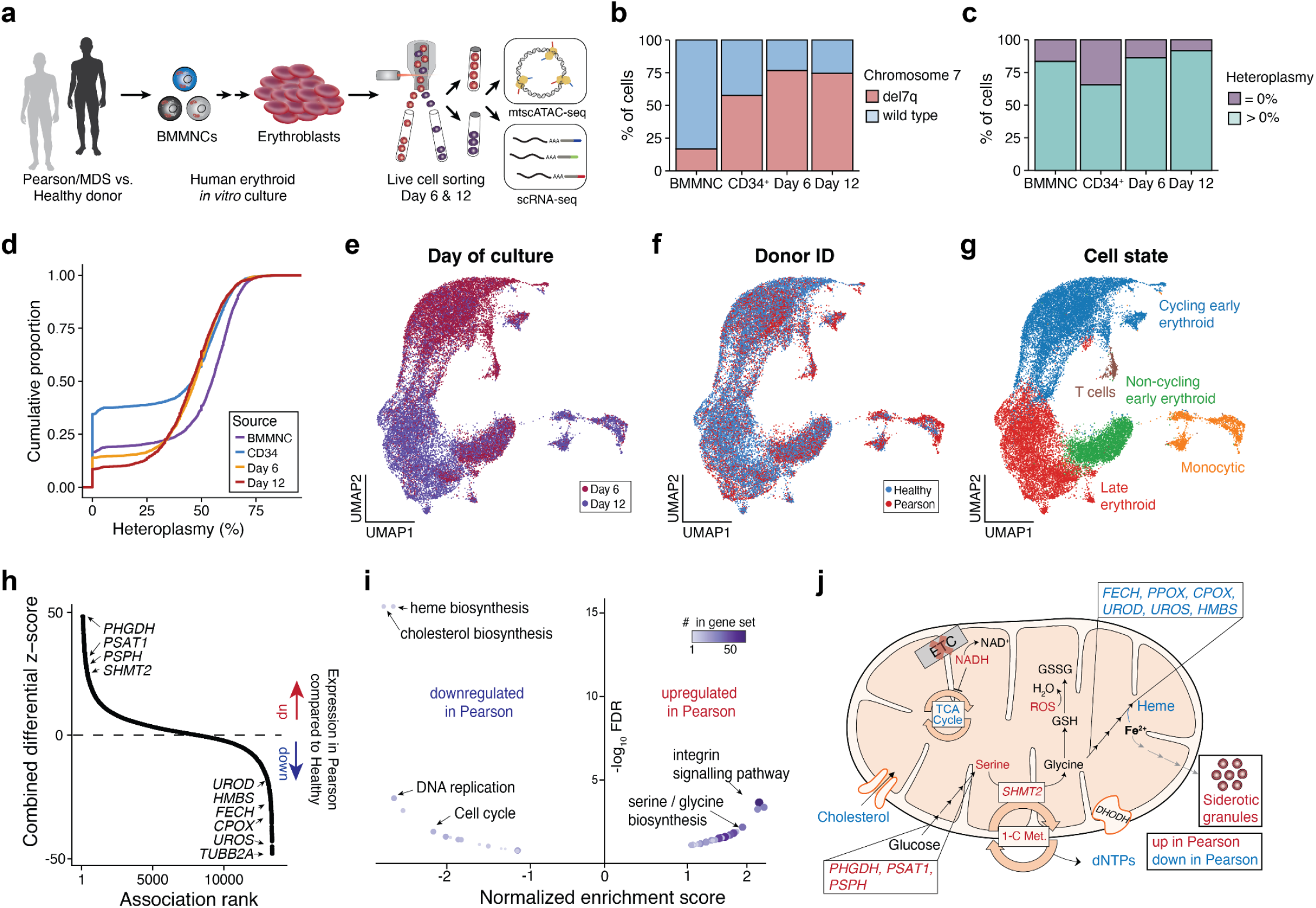
Altered *in vitro* erythroid differentiation and cellular states in Pearson syndrome. **(a)** Schematic of experimental design. Bone marrow mononuclear cells (BMMNCs) were derived from PT3 with PS/MDS and healthy controls and differentiated towards erythroblasts *in vitro*. Patient and healthy cells were harvested at day 6 and day 12 and jointly processed via mtscATAC-seq or scRNA-seq. **(b)** Stacked bar graph of cells annotated as wild type or del7q across indicated cell populations, including at day 6 and day 12 of *in vitro* culture. **(c)** Same as (b) but showing the proportion of cells with exactly 0% and >0% heteroplasmy of the mtDNA deletion as determined by mgatk-del. **(d)** Cumulative distribution graphs of mtDNA deletion heteroplasmy across the indicated four cell populations. **(e-g)** UMAP embedding of 28,783 high-quality cells profiled via scRNA-seq annotated by **(e)** day of culture collected, **(f)** healthy or disease state, and **(g)** annotated cell state/cluster. **(h)** Rank-sorted differentially expressed genes across erythroid populations. Select top genes overexpressed and down-regulated in PS are annotated. **(i)** Volcano plot of pathway enrichment analysis results via erythroid differential gene expression comparisons of the PS to healthy control cells. Select top pathways are annotated. **(j)** Schematic overview of altered (metabolic) genes and pathways in PS relative to the healthy stats. Genes and pathways upregulated in PS are shown in red and when downregulated shown in blue. Note: not all biochemical steps take place in mitochondria and the schematic has been simplified for illustrative purposes.

Next, we investigated the altered gene expression programs that may underlie the anemia phenotype in PS. To assess this, we performed unsupervised dimension reduction of 28,783 high-quality cells, which revealed a continuous trajectory of both PS and healthy donor cells undergoing erythroid differentiation as well as non-erythroid cells in discrete clusters (**Fig. 6e-g** and **Extended Data Fig. 6d; Methods**). We performed differential gene expression and pathway enrichment analyses comparing erythroid cells from the PS donor to the healthy control (**Fig. 6h,i; Methods**). Overall, the overlap of differentially expressed genes in both PBMCs and the *in vitro* cultured erythroid cells revealed no statistically significant enrichment of pathways (**Extended Data Fig. 6c; Methods**), and PS-specific genes overexpressed in PBMCs such as *CXCL14, C12orf54,* and *LINC01651* (**Fig. 3b**) were not expressed in erythroid cells, indicating multiple distinct and lineage-specific transcriptional changes in PS. Focusing on differentially expressed features in the erythroid culture, we observed genes of the serine and glycine biosynthesis pathway, in particular *PHGDH, PSAT1, PSPH,* and *SHMT2* to be significantly upregulated in PS erythroblasts (**Fig. 6h,i** and **Extended Data Fig. 6e**). Notably, serine metabolism has been reported to be altered in response to mitochondrial dysfunction and may aid to maintain cellular one-carbon availability to provide essential precursors for the synthesis of biomolecules such as purines, phospholipids, and the antioxidant glutathione (GSH), a scavenger of reactive oxygen species (ROS)^43–45^. Conversely, the heme biosynthesis pathway, including genes *UROS, CPOX, FECH, UROD, HMBS,*and *PPOX,* and cholesterol biosynthesis pathways were significantly downregulated in PS (**Fig. 6h,i** and **Extended Data Fig. 6**). In addition, we identified heme binding protein 2 *(HEBP2,* also known as *SOUL*), a gene associated with oxidative stress and mitochondrial permeabilization^46,47^, as a gene specifically expressed in PS erythroid cells (**Fig. 6g**). In total, our multi-omic analyses nominate numerous perturbed genes and pathways, the deregulation of which likely contribute to the characteristic anemia in PS (**Fig. 6j**), opening up multiple avenues for functional follow up and possible therapeutic interventions.

## Discussion

Advances in single-cell genomics have facilitated the comprehensive characterization of genomic features and their heterogeneity that define molecular phenotypes in health and disease states. Moreover, recently developed multi-omic approaches have started to provide complementary and orthogonal measurements and have tremendous potential to more holistically characterize the cellular circuits underlying perturbed cellular phenotypes in disease^48,49^. Here, we utilize these approaches to chart the genomic alterations across up to four modalities (i.e. transcriptome, accessible chromatin, mtDNA genotypes and nuclear chromosomal aberrations) resulting from large mtDNA deletions characteristic of PS, across tens of thousands of primary patient cells. In particular, we show how the development of mgatk-del in conjunction with mtscATAC-seq^9,10^ and ASAP-seq^25^ enables the identification and quantification of large mtDNA deletions in single cells together with concomitant readouts of cell state. This combination is of particular relevance given the unique features of mtDNA variants/deletions, which may be present at highly variable levels of heteroplasmy across a population of cells and their phenotypic effects being a function of heteroplasmy. In this regard, our advances will further facilitate the study of mtDNA mutations arising somatically in otherwise healthy individuals where these have been implicated to be a prominent contributor to in particular ageing phenotypes^1,2,50,51^. In light of recent reports noting the accumulation of mtDNA deletions to be more common in post-mitotic cells, including in single muscle fibers^52^ and neurons in Parkinson’s disease^53^, whereas SNVs appear more common in mitotic cells^54^, indicative of distinct evolutionary processes underlying these two classes of mutations, our framework now enables the comprehensive charting of these somatic mutations across a range of tissues.

In our application to PS, we consistently observed the depletion of pathogenic mtDNA deletions in CD8.TEM, is suggestive of purifying selection where the presence of pathogenic mtDNA does not appear to be compatible with the formation or maintenance of this cell state. In addition, we made analogous observations in both CD4+ and CD8+ RTEs as well CLPs derived from the bone marrow, indicating multiple modes of purifying selection and metabolic vulnerabilities at distinct stages of T cell maturation. These results are reminiscent of and refine our previous observation of purifying selection of the 3243A>G allele in T cells of adults with MELAS^10^. As the PS patients were pediatric (aged 4-7) this suggests the underlying selective processes related to pathogenic mtDNA to already occur early in life and also at different stages of lymphoid development. For example, we note that in peripheral blood, CD8.TEM, but no CD4+ T cell population, were predominantly purified of pathogenic mtDNA, corroborating distinct mitochondrial/metabolic demand following TCR-specific activation and conversion of CD8 T cells to effector/memory cells^55^. Such a model is consistent with the mitochondrial respiratory capacity to be a critical regulator of CD8 memory T cell development^56^. In contrast, in the bone marrow, CD4 and CD8 RTEs appeared to be equally affected, likely stemming from a purified lymphoid progenitor population (**Fig. 5**). Overall, our results provide novel insights into our emerging understanding of the effects of pathogenic mtDNA on the development of T cell states. Although additional studies will be required to elucidate the pathophysiological and metabolic defects underlying these observations in more detail, the systematic interrogation, including longitudinal profiling, of patients with distinct mtDNA disorders will shed further light on this multifaceted form of purifying selection.

More broadly, our observations surrounding the selection dynamics attributable to pathogenic mtDNA impose the question of their contribution to various clinical and immune-related phenotypes associated with primary mitochondrial disorders. Specifically, cohort studies have reported 42% of patients with mitochondrial disorders experience serious or recurrent infections, including 12% with sepsis; rates that are orders of magnitude more common than in the general population^15^. As such, we speculate that alterations in the T cell repertoire as well as cell state-specific metabolic requirements compromise eliciting an effective immune response during infections. Notably, patients with mitochondrial disorders also have a reported >4x incidence of primary tumors^14^. While this increased prevalence has previously been attributed to alterations in metabolism which may intrinsically favor the evolution of cancer cells^57^, we suggest that in addition metabolic alterations within the pool of circulating anti-tumor T cells may further diminish mounting an effective anti-tumor response.

In a case of PS with MDS, we further examined the complex interplay between the pathogenic mtDNA deletion and a somatically acquired del7q that is associated with MDS (**Fig. 4** and **5**). Interestingly, the del7q is restricted to cells carrying the mtDNA deletion, suggesting that the del7q allele arose in a mitochondrially/metabolically dysfunctional background. Conceivably, the resulting competitive advantage of the del7q is more pronounced than in mtDNA wildtype cells. Moreover, the distinct distributions of cells across the genomics landscapes and carrying the del7q and mtDNA deletion is notable, with their study being uniquely enabled by approaches such as mtscATAC-seq and ASAP-seq. Moreover, the del7q appears to lead to an expansion of myeloid cells and is largely absent in T cells. CLPs also appear to be relatively depleted of pathogenic mtDNA and the phenotypic HSC pool also appears significantly diminished though we cannot fully disentangle to what extent this may be a consequence of the MDS or PS or in fact both. In this regard, we believe that a refined understanding of lineage commitment and differentiation, also incorporating lineage and clonal tracing readouts will be essential to the study of such pathologic contexts in human hematopoiesis. Here, it is noteworthy that somatic mtDNA mutations occurring early in life in the investigated pediatric patients show the potential to further facilitate such efforts.

In an *in vitro* model, we further assessed genomic and mtDNA features during erythroid differentiation to enhance our understanding of the characteristic sideroblastic anemia in PS (**Fig. 6**). Collectively, the pathway enrichment and differential gene expression results provide an overview of the alterations that may cooperatively contribute to the underlying erythroid defect (**Fig. 6j**). Specifically, cholesterol biosynthesis, heme biosynthesis, and serine/glycine metabolism appear significantly altered in PS, with defects in cholesterol synthesis being associated with impaired red blood cell differentiation^58^. In light of the increased NADH:NAD+ ratios and ROS levels associated with mtDNA-related diseases as a consequence of the impaired electron transport chain^59–61^, we suggest that serine/glycine biosynthesis is upregulated in PS cells, as glycine serves as a precursor for glutathione generation (GSH, a ROS scavenger) as well as for the remodeling of one-carbon metabolism to maintain production of DNA and other critical components of the cell in states of hypoxia or mitochondrial dysfunction^43,62^. In this regard, mitochondrial one-carbon metabolism appears less sensitive to product inhibition by increased NADH:NAD+ ratios^45^. Elevated NADH:NAD+ ratios and ROS levels may further contribute to the downregulation of heme biosynthesis^63^ although heme, ultimately with iron, is essential for the adequate production of hemoglobin during red blood cell generation^64^. As glycine is also a necessary precursor of heme biosynthesis, it may however be rechanneled to synthesize one carbon precursors for DNA replication in normally highly proliferative erythroblasts and/or GSH to scavenge increased ROS levels resulting from mitochondrial dysfunction. Consequently, heme biosynthesis genes may be downregulated and excess iron accumulates, leading to granular depositions and the formation of the characteristic sideroblastic cells in PS. Importantly, excess iron may further contribute to the formation of reactive oxygen species^65,66^, thereby further exacerbating the need for GSH.

In total, our multi-omic methods revealed unique genomic alterations in response to pathogenic mtDNA in distinct cellular compartments, such as PBMCs and differentiating erythroblasts. While mitochondria are ubiquitous, they nevertheless fulfill distinct roles as a function of cell type and cell state. This emphasizes the need to ideally study patient-derived cellular specimens where possible to fully capture alterations resulting from mitochondrial dysfunction attributable to germline or somatic mtDNA mutations. In this light, we demonstrate how comprehensive single-cell multi-omic approaches provide biologically important insights into the molecular alterations of primary mitochondrial defects as well as a rich resource for functional follow-up studies. Together with orthogonal metabolic readouts and their adaptations to single cells^67,68^, we envision to further improve our understanding of how selective pressures in the mitochondrial genome contribute to complex human phenotypes.

## EXTENDED DATA– FIGURES AND LEGENDS

**Extended Data Fig. 1.**
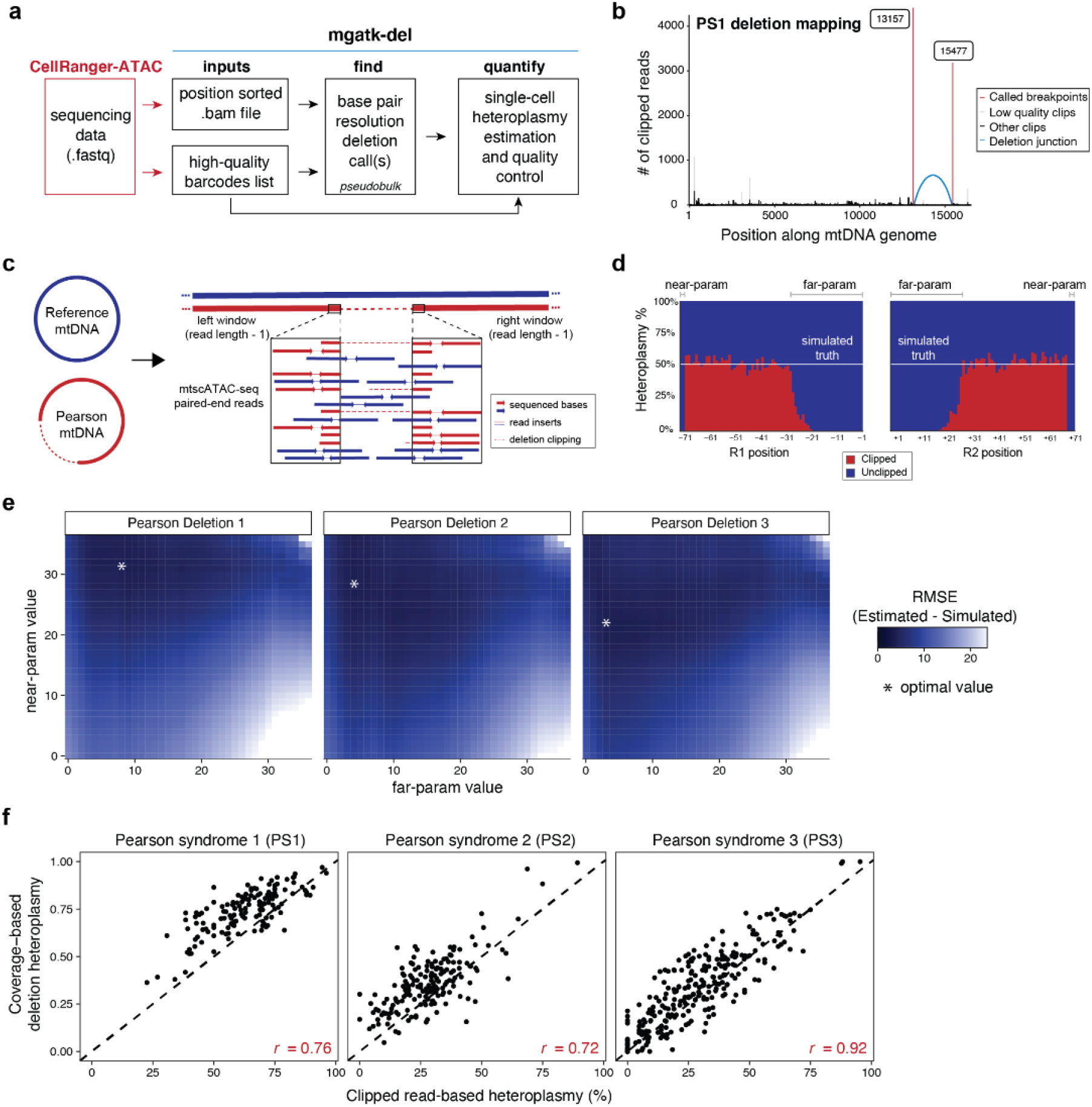
Supporting information for mtDNA deletion calling and heteroplasmy estimation using mgatk-del. **(a)** Schematic of mgatk-del pipeline, which utilizes the outputs of CellRanger-ATAC. Two critical steps of base-resolution deletion calling (‘find’) and estimation of single-cell heteroplasmy (‘quantify’) are illustrated. **(b)** Output of mgatk-del ‘find’ for Pearson syndrome deletion 1 (PS1). The red vertical lines represent the called deletion breakpoints, which are spanned by the blue arc signifying the number of reads where the regions were joined via a secondary alignment (‘SA’ tag in .bam file). **(c)** Schematic of simulation framework. Synthetic cells with known heteroplasmy were generated via mixtures of reference and PS mtDNA for all previously reported deletions. Colors represent the originating genome from the simulation as shown on the left. **(d)** Summary of results from a 50% mix for a mtDNA deletion showing the estimated heteroplasmy as estimated from the ratio of clipped to unclipped reads. The critical parameters ‘near-param’ and ‘far-param’ define the number of bases that are discarded on the read when estimating the overall heteroplasmy per cell. **(e)** Results of exhaustive simulation for three mtDNA deletions used in the cell mixing experiment. The minimum value of the root mean squared error (RMSE) of the estimated and true heteroplasmy is noted with an asterisk over the grid search. **(f)** Single-cell correlation of clipped (Fig. 1d) versus coverage-based (Fig. 1e) heteroplasmy estimates for valid deletions per indicated deletion/donor. The Pearson correlation for the three deletions for both modes of heteroplasmy estimation is indicated.

**Extended Data Fig. 2.**
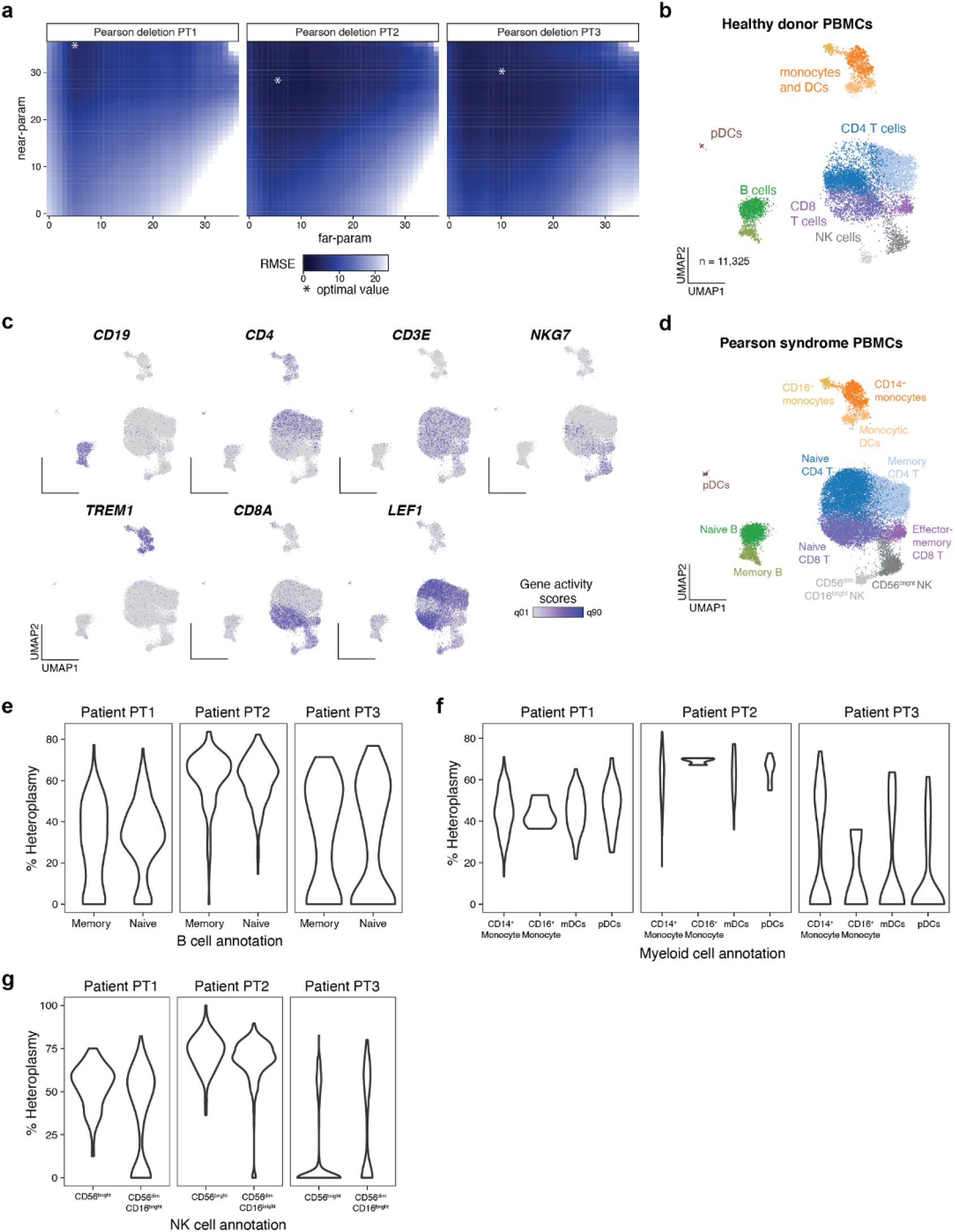
Supporting information for Pearson syndrome PBMC mtscATAC-seq analyses. **(a)** Result of mgatk-del hyperparameter optimization via a simulation framework. The minimum value of the root mean squared error (RMSE) of the estimated and true heteroplasmy is noted with an asterisk over the grid search. **(b)** UMAP embedding of healthy donor control PBMCs. Major cell clusters/types are indicated. n is the number of cells assayed. **(c)** Embedding as in (b) and visualization of select marker genes used to support cell type annotation based on chromatin accessibility analysis. Shown are the normalized gene activity scores computed by Signac, and the color is scaled to the indicated quantiles on the color bar. **(d)** UMAP and annotation of identified cell types in the PS PBMC samples; compare to Fig. 2d,e. **(e-g)** Violin plots of **(e)** B cell, **(f)** myeloid cell, and **(g)** natural killer cell subsets indicating the distribution of heteroplasmy of each respective mtDNA deletion for all three patients.

**Extended Data Fig. 3.**
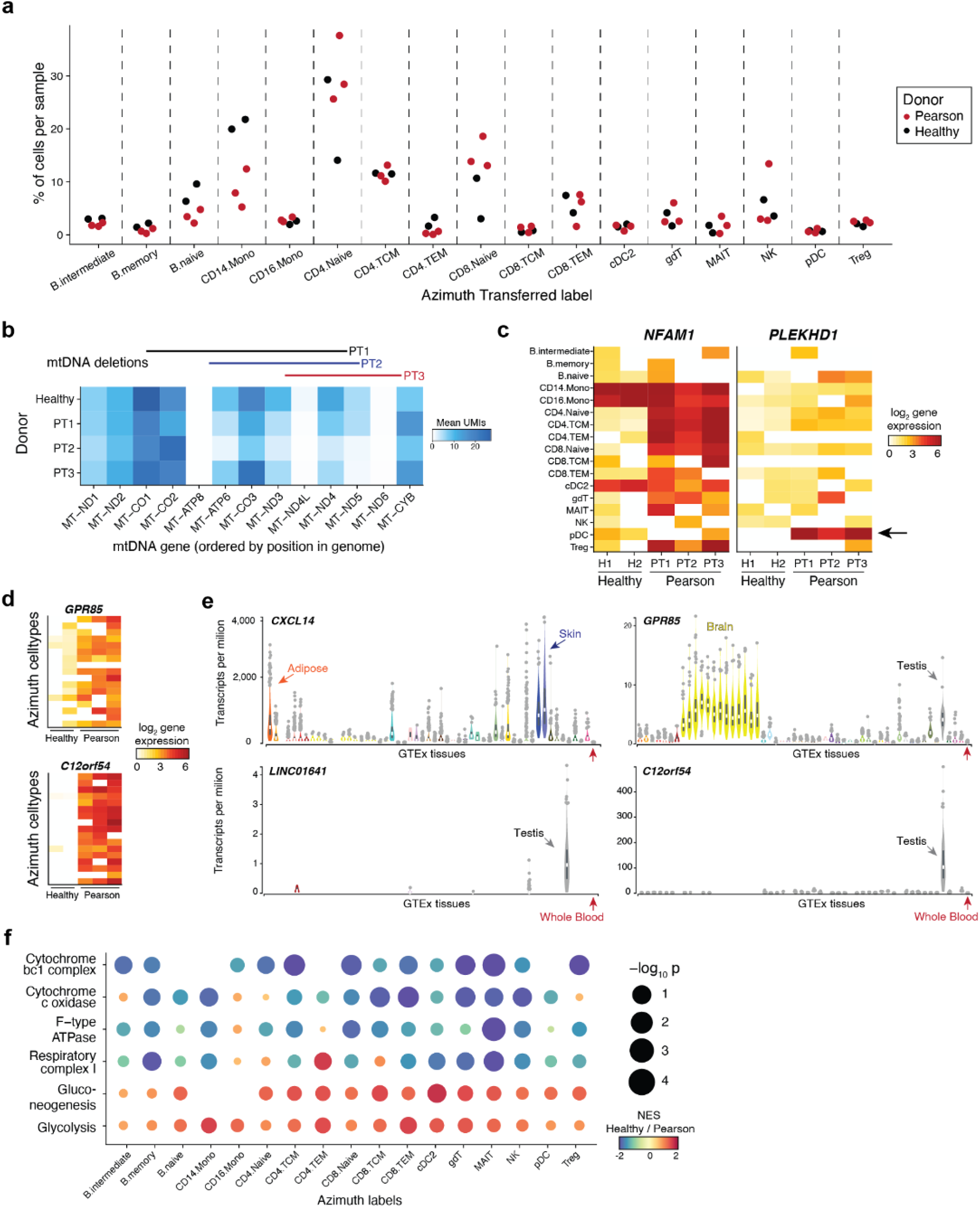
Supporting information for Pearson syndrome PBMC scRNA-seq analyses. **(a)** Proportions of 17 major PBMC types after reference projection per patient/ healthy donor. **(b)** Summary of mean gene expression for indicated genes encoded in mtDNA ordered by position in the contig. Bars above indicate the respective deletions for the three PS patients. **(c,d)** Heatmap of pseudobulk gene expression across PBMC types for indicated genes. (c) *NFAM1* and *PLEKHD1* are specifically up-regulated in PS compared to healthy controls in a subset of cells, e.g. CD4 T cells. (d) Genes *GPR85* and *C12orf54* are upregulated in all cell types in PS compared to a healthy control. **(e)** Summary of GTEx derived gene expression across tissues of select differentially expressed genes in PS PBMCs. Whole blood (the closest comparator to PBMCs) is highlighted with a red arrow. **(f)** Gene set enrichment analysis for indicated metabolic pathways across cell types. Following the indicated color gradient, more red indicates genes to be upregulated in a pathway for healthy donors relative to PS cells, whereas more purple indicates genes in a pathway that are upregulated in PS compared to healthy donor cells.

**Extended Data Fig. 4.**
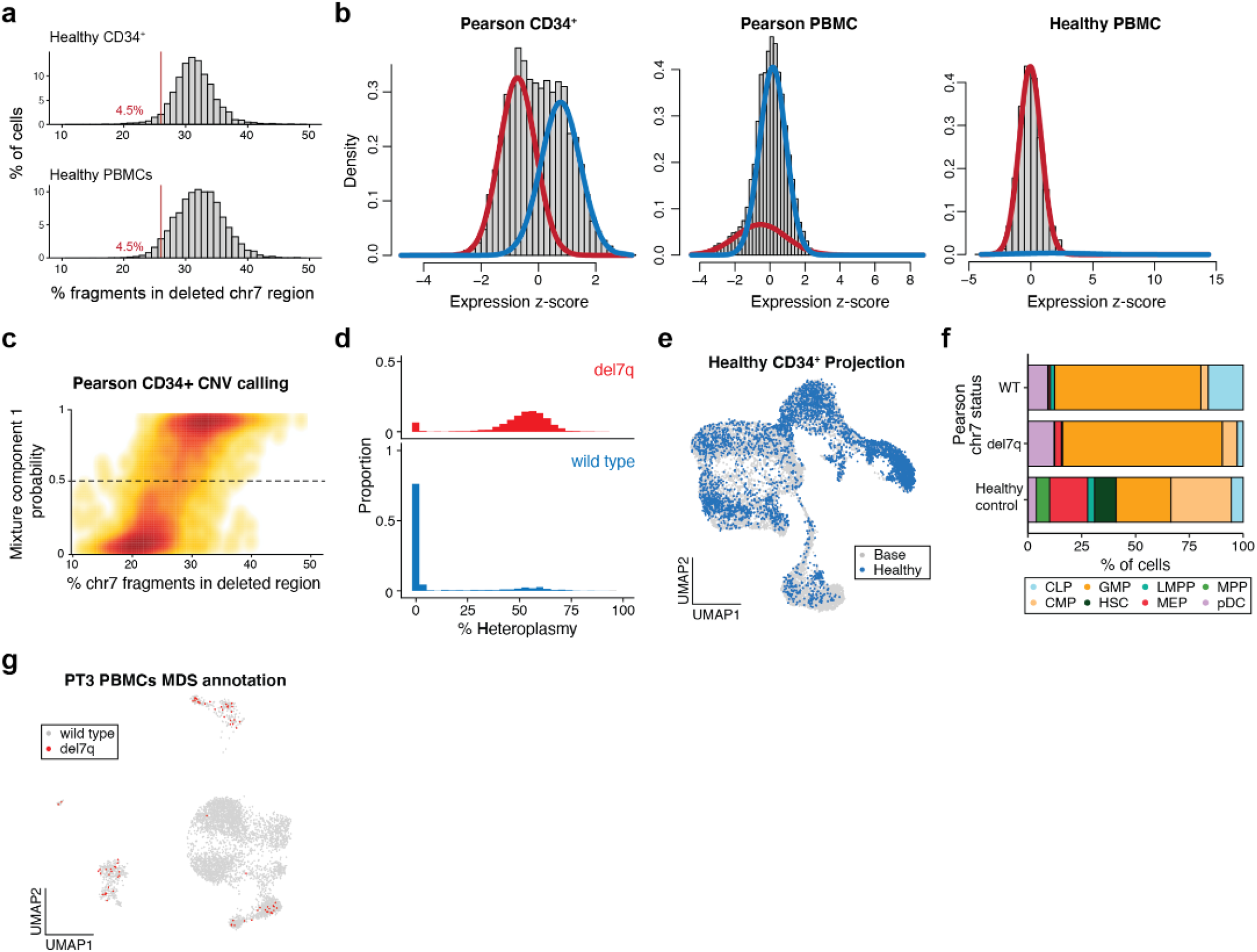
Supporting information for del7q calling and CD34+ mtscATAC-seq analyses. **(a)** Summary of 7q fragment abundances in healthy CD34+ and PBMC mtscATAC-seq samples^9^; compare to Fig. 4b with the same cutoff. **(b)** Result of Gaussian mixture model applied to indicated samples. The red trace indicates the first mixture component estimated (lower mean) whereas the blue trace represents the second component with higher mean. The healthy PBMC sample does not contain a chromosome alteration. **(c)** Graphical density of cells from mixture model (y-axis) and from crude fragment abundance (x-axis; see Fig. 4b). The dotted line indicates the cutoff for wild type and del7q annotations. **(d)** Histograms of mtDNA deletion heteroplasmy proportions (%) stratified on del7q status. **(e)** Projection of a healthy control CD34+ mtscATAC-seq sample^9^ onto the reference embedding as shown in Fig. 4d. **(f)** Stacked bar plots of cell type proportions for projected cell types from PT3 with PS/MDS stratified by del7q status (MDS for positive and wild type for negative) and a healthy control donor. **(g)** Annotation of del7q status in PBMCs, which is primarily identified in myeloid, NK, and B-cell populations; see Fig. 2D for cluster annotations.

**Extended Data Fig. 5.**
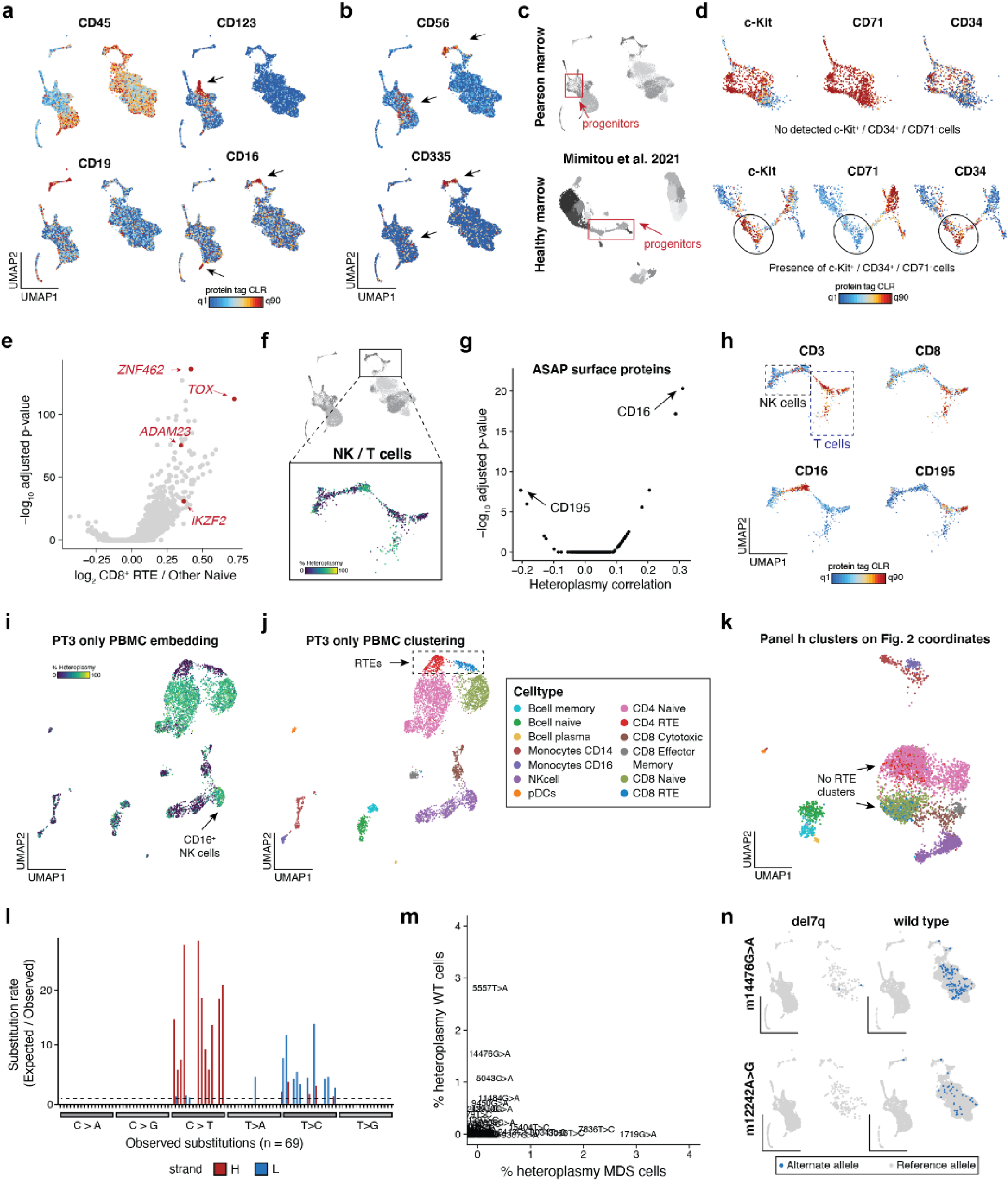
Supporting analyses for PT3 bone marrow mononuclear cell ASAP-seq dataset. **(a)** Projection of select protein-derived antibody tag abundances for indicated proteins. Select arrows indicate populations positive for the respective marker. See Fig. 5c for cluster annotations. **(b)** Projection of protein surface markers CD56 (NK cells and MDS-associated cells) and CD335 (only NK cells) with arrows indicating the two cell populations. **(c)** UMAP of ASAP-seq processed bone marrow mononuclear cells from a PS (top) and a healthy control^25^ (bottom) with hematopoietic stem and progenitor cells (‘progenitors’) indicated in the red boxes. **(d)** Projection of protein tags within the boxed progenitor populations as in (c), contrasting the presence of only CD71+ cells among CD34+/c-Kit+ cells in PS as compared to the healthy control. We note that for the PS sample, CD34 tag complexity is lower, as these cells were also co-stained with a fluorophore-conjugated antibody to enrich the CD34+ cell population via sorting. **(e)** Volcano plot showing differential gene activity scores for CD8 recent thymic emigrants (RTEs) compared to other CD8 naive T cells. Annotated genes in red represent known marker genes for RTEs. **(f)** Zoom (top) and mtDNA deletion heteroplasmy (bottom) in differentiated CD8 T and NK cells from the BMMNC populations. **(g)** Volcano plot illustrating the association between protein levels and mtDNA deletion heteroplasmy in single cells. The two top proteins in either effect direction are highlighted. **(h)** Projection of cell state surface markers (CD3, CD8) and top antibody tags (CD16, CD195) as determined in (g). **(i,j)** Reclustering and UMAP depiction of PT3 PBMC mtscATAC-seq data identifies **(i)** low heteroplasmy and **(j)** recent thymic emigrants (RTEs). Cell type annotations as indicated. **(k)** The same cluster annotations (colored) from panel (j) are used on the coordinates from the integrated UMAP depiction of all patients as in Fig. 2. **(l)** Substitution rate (observed over expected) of mgatk identified heteroplasmic mutations (y-axis) in each class of mononucleotide and trinucleotide change resolved by the heavy (H) and light (L) strands of the mitochondrial genome. **(m)** Scatter plot of 69 somatic mtDNA variants identified in panel (l) stratified based on cells annotated as del7q (x-axis) and wild type for chr7 (y-axis). **(n)** Projection of wild type (diploid chr7)-enriched somatic mtDNA mutations 14476G>A (50%) and 12242A>G (25%). Threshold for + cell heteroplasmy is based on the empirical density of each variant with the threshold being indicated in parentheses for each variant.

**Extended Data Fig. 6.**
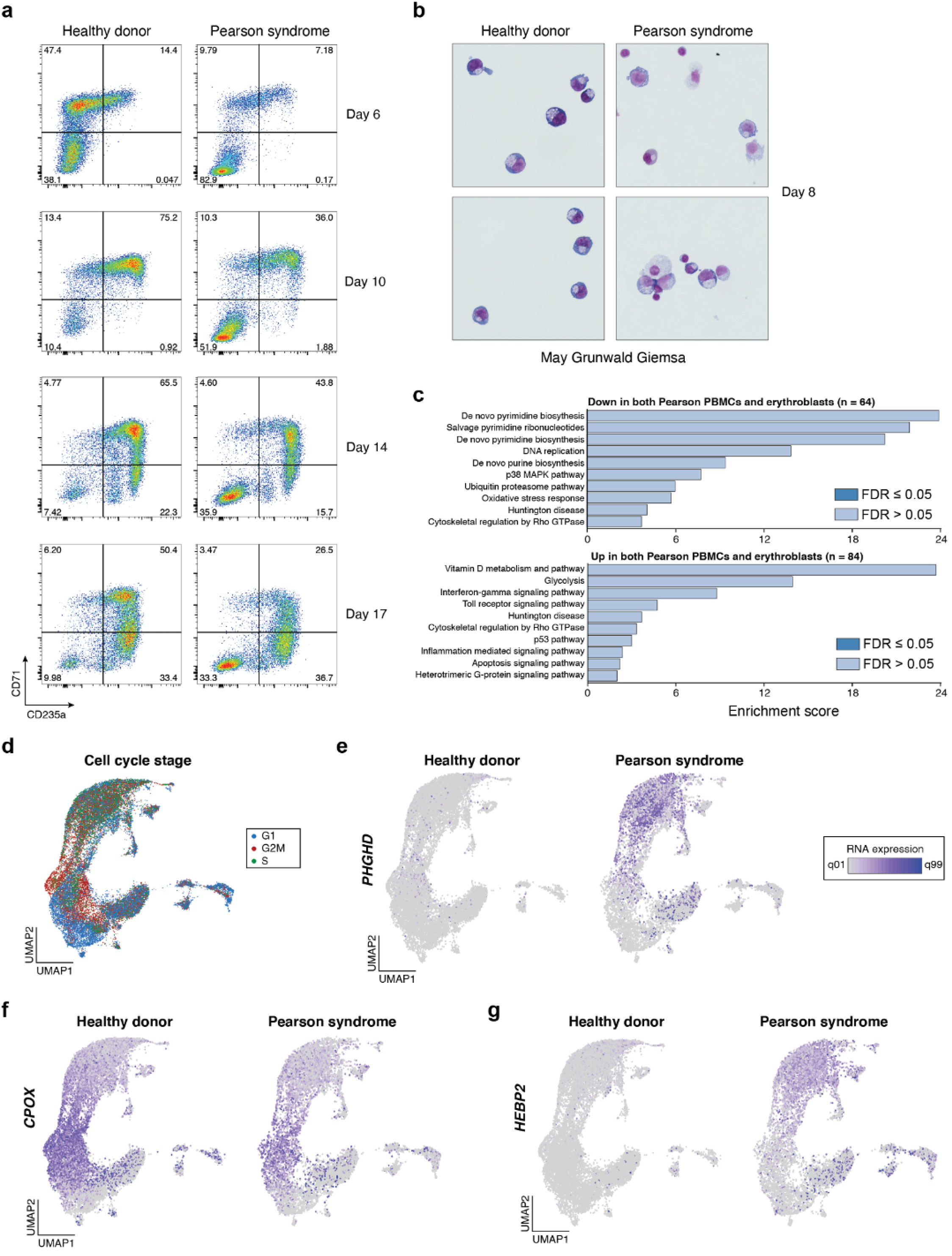
Supporting information for *in vitro* erythroid differentiation experiments. **(a)** Flow cytometry plots showing the distribution of CD71 and CD235a expression for *in vitro* differentiated healthy control and PS cells at indicated days of culture. **(b)** MayGrunwald Giemsa stained cytospins of *in vitro* differentiated healthy control and PS cells at day 8 of culture. **(c)** Pathway analysis of top up- and downregulated genes in both PS PBMCs and *in vitro* differentiated erythroblast scRNA-seq datasets. **(d)** UMAP of scRNA-seq data colored by predicted cell cycle state. Cluster annotations as in Fig. 6E-G. **(e-g)** Projection of gene expression of selected differentially expressed genes between PS and healthy control erythroblasts, including **(e)** *PHGDH,* **(f)** *CPOX,* and **(g)** *HEBP2*. Gene expression coloring is scaled for all plots between the first and 99th quantile per gene.

## ACKNOWLEDGEMENTS

We are deeply grateful to the patients and families who made this work possible. We are grateful to members of the Agarwal, Regev, Sankaran, and Ludwig laboratories for their helpful discussions. We thank Kar-Tong Tan and Matthew Meyerson for input concerning the 7q deletion. We thank Akiko Shimamura and the Boston Children’s Hospital Bone Marrow Failure and Myelodysplastic Syndrome program for their support in this study (supported by NIH grant RC2DK122533). We acknowledge support from the Broad Institute and the Whitehead Institute Flow Cytometry core facilities. CAL is supported by a Stanford Science Fellowship, a Parker Institute of Cancer Immunotherapy Scholarship, and NIH K99 HG012076. This research was supported by a BroadIgnite Award (LSL), a Google Cloud seed grant (CAL), by the United Mitochondrial Disease Foundation and the North American Mitochondrial Disease Registry NIH U54 (SP), a Lloyd J. Old STAR Award from the Cancer Research Institute (ATS), an ASH Scholar Award from the American Society of Hematology (ATS), R01 DK103794 (VGS), and R33 HL120791 (VGS), a gift from the Lodish Family to Boston Children’s Hospital (VGS), the New York Stem Cell Foundation (NYSCF, VGS), the Howard Hughes Medical Institute and Klarman Cell Observatory (AR), the Champ Foundation (SA) and the Associazione Luigi Comini Onlus (SA). YHH is supported by a PhD fellowship from the Hector Fellow Academy. PK and LN receive support as associate members of the Hector Fellow Academy. VGS is a NYSCF-Robertson Investigator. SA is supported by R01 DK107716 and R33 HL154133. LSL is supported by an Emmy Noether fellowship by the German Research Foundation (DFG, LU 2336/2-1), a Longevity Impetus grant, and a Hector Research Career Development Award by the Hector Fellow Academy. CAL, LSL, and ATS are supported by 1UM1HG012076-01.

## AUTHOR CONTRIBUTIONS

CAL, AR, VGS, SA, and LSL conceived and designed the project. LSL led, designed, and performed experiments with FAB, YHH, PK, LN, YY, and assistance from SMD, WL, and CM. EM and PS contributed reagents and advised on the ASAP-seq experiments. LO, JS, and CE contributed healthy pediatric control samples and clinical perspectives. CAL developed the mgatk-del software and led all analyses. CAL, KG, and JMV developed the mtDNA simulation framework. FB, YHH, PK, LN, SDP, JAG, EF, RM, PM, SuP, AK, SA, and LSL contributed to data interpretation. SMD, SDP, MDF, AK, ATS, AR, VGS, SA, and LSL supervised various aspects of this work. CAL and LSL wrote the manuscript with input from all authors.

## METHODS

### Cell lines and cell culture

Biological samples for cell lines were procured under protocols approved by the Institutional Review Board at Boston Children’s Hospital, after written informed consent in accordance with the Declaration of Helsinki. Pearson syndrome fibroblasts were derived from patient bone marrow (PS1 and PS2) or skin (PS3). Control fibroblasts were derived from healthy donors’ skin (Control1 and Control2). All fibroblasts were grown in DMEM containing 15% FBS, L-glutamine, non-essential amino acid and penicillin/streptomycin. Cells were incubated at 37°C with 5% CO_2_.

### Healthy donor and patient samples

Primary human peripheral blood and bone marrow samples were collected under Institutional Review Board-approved protocols and written informed consent for genomic sequencing. Primary hematopoietic samples were collected from three previously diagnosed patients, including a 7-year-old male with PS / Kearns Sayre syndrome (“PT1”), a 4-year-old female with PS (“PT2”), and a 4-year-old male with PS and deletion 7q (del7q) myelodysplastic syndrome (MDS; “PT3”). Peripheral blood and bone marrow mononuclear cells were isolated using Ficoll Paque Plus solution and density gradient centrifugation using SepMate tubes (StemCell Technologies). All samples were stored in vapor-phase liquid nitrogen after cryopreservation with 10% dimethyl sulfoxide until analysis. Healthy donor bone marrow mononuclear cells were obtained from StemCell Technologies. Healthy adult CD34^+^ hematopoietic stem and progenitor cells (HSPCs) were obtained from the Fred Hutchinson Hematopoietic Cell Processing and Repository (Seattle, USA). The CD34+ samples were de-identified and approval for use of these samples for research purposes was provided by the Institutional Review Board and Biosafety Committees at Boston Children’s Hospital.

### Human erythroid *in vitro* cell culture

Bone marrow mononuclear cells or CD34+ HSPCs from healthy donors or PT3 were differentiated into mature erythroid cells utilizing a three-phase culture protocol^69,70^. Cells used for scRNA-seq and mtscATAC-seq experiments were derived from two independent cultures using PT3 cells, but two different healthy control donors were used for each culture. In phase 1 (day 0 – 7), cells were cultured at a density of 10^5^ - 10^6^ cells/ml in IMDM supplemented with 2% human AB plasma, 3% human AB serum, 1% penicillin/streptomycin, 3 IU/ml heparin, 10 μg/ml insulin, 200 μg/ml holo-transferrin, 1 IU erythropoietin (Epo), 10 ng/ml stem cell factor (SCF) and 1 ng/ml IL-3. In phase 2 (day 7 – 12), IL-3 was omitted from the medium. In phase 3 (day 12 – 18), cells were cultured at a density of 1×10^6^ cells/ml, with both IL-3 and SCF omitted from the medium and the holo-transferrin concentration was increased to 1 mg/ml. Cells were cultured at 37°C and 5% CO_2_.

### Flow cytometry analysis and sorting

For flow cytometry analysis and sorting, cells were washed in FACS buffer (1% FBS in PBS) before antibody staining. *In vitro* cultured primary erythroid cells were stained using 1:50 APC-conjugated CD235a (Glycophorin A, clone HIR2, 50-153-69, eBioscience) and 1:50 FITC-conjugated CD71 (clone OKT9, 14-0719-82, eBioscience) for 15 min on ice. PT3 bone marrow-derived CD34+ cells were stained using 1:40 APC-conjugated CD34 (clone 581, 343509, BioLegend). For mtscATAC-seq and ASAP-seq experiments residual granulocytes were excluded by staining of cells using 1:50 PE-conjugated CD66b (clone G10F5, 305102, BioLegend). For live/ dead cell discrimination Sytox Blue was used at a 1:1000 dilution according to the manufacturer’s instructions (Thermo Fisher, S34857). FACS analysis was conducted on a BD Bioscience Fortessa flow cytometer at the Whitehead Institute Flow Cytometry core. Data was analyzed using FlowJo software v10.4.2. Cell sorting was conducted using the Sony SH800 sorter with a 100 μm chip at the Broad Institute Flow Cytometry Facility.

### Mitochondrial single-cell ATAC-seq (mtscATAC-seq)

MtscATAC-seq libraries were generated using the 10x Chromium Controller and the Chromium Single Cell ATAC Library & Gel Bead Kit (#1000111) according to the manufacturer’s instructions (CG000169-Rev C; CG000168-Rev B) as outlined below and previously described to increase mtDNA yield and genome coverage^9^. Briefly, 1.5 ml or 2 ml DNA LoBind tubes (Eppendorf) were used to wash cells in PBS and downstream processing steps. After washing cells were fixed in 0.1 or 1% formaldehyde (FA; ThermoFisher #28906) in PBS for 10 min at RT, quenched with glycine solution to a final concentration of 0.125 M before washing cells twice in PBS via centrifugation at 400 g, 5 min, 4°C. Cells were subsequently treated with lysis buffer (10mM Tris-HCL pH 7.4, 10mM NaCl, 3mM MgCl_2_, 0.1% NP40, 1% BSA) for 3 min for primary cells and 5 min for cell lines on ice, followed by adding 1 ml of chilled wash buffer and inversion (10mM Tris-HCL pH 7.4, 10mM NaCl, 3mM MgCl_2_, 1% BSA) before centrifugation at 500 g, 5 min, 4°C. The supernatant was discarded and cells were diluted in 1x Diluted Nuclei buffer (10x Genomics) before counting using Trypan Blue and a Countess II FL Automated Cell Counter. If large cell clumps were observed a 40 μm Flowmi cell strainer was used prior to processing cells according to the Chromium Single Cell ATAC Solution user guide with no additional modifications. Briefly, after tagmentation, the cells were loaded on a Chromium controller Single-Cell Instrument to generate single-cell Gel Bead-In-Emulsions (GEMs) followed by linear PCR as described in the protocol using a C1000 Touch Thermal cycler with 96-Deep Well Reaction Module (BioRad). After breaking the GEMs, the barcoded tagmented DNA was purified and further amplified to enable sample indexing and enrichment of scATAC-seq libraries. The final libraries were quantified using a Qubit dsDNA HS Assay kit (Invitrogen) and a High Sensitivity DNA chip run on a Bioanalyzer 2100 system (Agilent).

### ATAC with Selected Antigen profiling by sequencing (ASAP-seq)

PT3-derived bone marrow mononuclear cells were stained with a 242 TSA-conjugated antibody panel as previously described^25^. To enable flow cytometry based enrichment of CD34+ cells, the sample was co-stained using an APC-conjugated CD34 (clone 581, 343509, BioLegend) to sort live CD66b-CD34+ and otherwise CD66b-bone marrow mononuclear cells, which were then pooled after sorting and processed for ASAP-seq as previously described^25^ and outlined online: https://cite-seq.com/asapseq/. Briefly, following sorting, cells were fixed in 1% formaldehyde and processed as described for the mtscATAC-seq workflow described above, with the modification that during the barcoding reaction, 0.5μl of 1μM bridge oligo A (BOA for TSA) was added to the barcoding mix. Sequence of BOA: TCGTCGGCAGCGTCAGATGTGTATAAGAGACAGNNNNNNNNNVTTTTTTTTTTTTTTTTTTTTTTTTTTTTTT/ 3InvdT/. For GEM incubation the standard thermocycler conditions were used as described by 10x Genomics for scATAC-seq. Silane bead elution and SPRI cleanup steps were modified as described to generate the indexed protein tag library^25^. The final libraries were quantified using a Qubit dsDNA HS Assay kit (Invitrogen) and a High Sensitivity DNA chip run on a Bioanalyzer 2100 system (Agilent).

### Single-cell RNA-seq

ScRNA-seq libraries were generated using the 10x Chromium Controller and the Chromium Single Cell 3’ Library Construction Kit v2 according to the manufacturer’s instructions. Briefly, the suspended cells were loaded on a Chromium controller Single-Cell Instrument to generate single-cell Gel Bead-In-Emulsions (GEMs) followed by reverse transcription and sample indexing using a C1000 Touch Thermal cycler with 96-Deep Well Reaction Module (BioRad). After breaking the GEMs, the barcoded cDNA was purified and amplified, followed by fragmenting, A-tailing and ligation with adaptors. Finally, PCR amplification was performed to enable sample indexing and enrichment of scRNA-Seq libraries. The final libraries were quantified using a Qubit dsDNA HS Assay kit (Invitrogen) and a High Sensitivity DNA chip run on a Bioanalyzer 2100 system (Agilent).

### Sequencing

All libraries were sequenced using Nextseq High Output Cartridge kits and a Nextseq 550 sequencer (Illumina). 10x scATAC-seq and ASAP-seq libraries were sequenced with paired-end reads (2 × 72 cycles). 10x 3’ scRNA-seq libraries were sequenced as recommended by the manufacturer.

### mtDNA deletion calling and heteroplasmy estimation in single cells

Though large mtDNA deletions have been well-documented in a variety of next-generation sequencing datasets, we observed that coordinates associated with deletions (e.g. the ‘common’ deletion) may be incorrect at base pair resolution. These differences are primarily due to differences in the coordinates of the mitochondrial reference genome and variation in the results of sequencing read alignment tools, particularly near homomorphic sequences at deletion junctions. Thus, we recommend utilizing the sequencing data from the particular sequencing experiment to identify the deletion junction within the primary sequencing data that is being analyzed. As part of our software solution, mgatk-del in “find” mode takes a .bam file and compiles a list of key summary statistics, including the number of clipped reads per position, secondary alignment bases, and overall coverage, in order to identify deletions. The outputs of mgatk-del find include plots (e.g. **Extended Data Fig. 1B**) and tables that facilitate identifying the specific base pairs associated with the mtDNA deletions.

After the precise break points have been determined, single-cell mtDNA deletion heteroplasmy can be estimated using the second step in the mgatk-del pipeline. Here, PCR-deduplicated single-cell bam files (emitted as intermediate output in the standard mgatk pipeline) serve as the primary input, yielding an estimation of mtDNA deletion heteroplasmy per user-specified deletion. This metric is determined by using the ratio of reads overlapping the deletion junction that either support (via clip) or provide no evidence of the deletion (contiguous alignment over the read window as depicted in **Fig. 1c,d**). Notably, each paired-end read contributes only once to the heteroplasmy metric. As a comparison, coverage based heteroplasmy (**Fig. 1e**) was estimated via one minus the ratio of mean per-base coverage within the deleted region over the mean per-base coverage outside the deleted region. For negative values (when the within deleted region coverage exceeded that of the outside region), values were adjusted to 0% heteroplasmy for display purposes (coverage based heteroplasmy was not used in any downstream analyses).

To generate simulated sequencing datasets and to benchmark this approach, we used the wgsim tool from within samtools^71^ to generate paired-end reads each of length 72 bp (same length as for our mtscATAC-seq data), 50 bp, or 100 bp, which represent common sequencing configurations. Sequencing reads were simulated from either the revised Cambridge reference sequence (rCRS) or a synthetic mtDNA chromosome that encoded the specified deletion. Simulated sequencing reads were then aligned to the masked reference genome used by CellRanger-ATAC, and the resulting aligned reads from the .bam files were mixed in specific ratios (10 mixtures per deletion) to specify the true heteroplasmy for the given simulation. The estimated heteroplasmy was computed running the function utilized in mgatk-del with the search space of possible values in the near-param and far-param. Then the root-mean squared error (RMSE) was computed based on the difference. In total, 22 mtDNA deletions were considered, which represented a curated list of the six deletions in our study and 16 additional deletions that were curated from MITOMAP^72^. The default parameters in mgatk-del (near-param: 9; far-param: 24) represent values that performed consistently well across a variety of deletions and read lengths. We suggest that mgatk-del can produce reasonable single-cell heteroplasmy estimations from the default parameters. Specifically, we observed a mean 0.93% RMSE difference between default and optimal hyper-parameter values across our six PS mtDNA deletions in the cell lines and primary cells. Thus, we suggest that a grid-search to determine optimal hyper-parameters for accurate heteroplasmy estimation may be useful but typically unnecessary for new datasets.

### scATAC-seq analyses

Raw sequencing data was demultiplexed using CellRanger-ATAC mkfastq. Demultiplexed sequencing reads for all libraries were aligned to the mtDNA blacklist modified^9^ hg19 reference genome using CellRanger-ATAC count v1.2. Deletions in mtDNA were identified per patient library and heteroplasmy was quantified using the exact breakpoints as discussed in the previous section. Downstream analyses of the three PS donors and one healthy donor previously profiled with mtscATAC-seq^9^ were performed after identifying cells with a minimum depth of 10x on mtDNA, 1000 ATAC fragments passing filters, and 45% of fragments in accessibility peaks from an aggregated peak set. Latent semantic indexing (LSI) was performed and the 2-30 components were adjusted for donor effects using harmony before producing a two-dimensional embedding and clustering using the harmony components^73^. Gene activity scores were computed and normalized using the Signac workflow, and heteroplasmy-gene correlations were computed using per-donor scaled heteroplasmy values for the T cell clusters (determined by CD3 accessibility). Transcription factor activity scores were computed via chromVAR using the Signac wrapper functions.

### scRNA-seq analyses

PS cell-derived 10x 3’ scRNA-seq sequencing libraries were demultiplexed and aligned to the hg38 reference with CellRanger v3.0.2. Healthy PBMC datasets were augmented from the public resource of 10x single-cell gene expression. Raw sequencing reads from two libraries (pbmc4k; pbmc8k) of 10x v2 chemistry were downloaded and reprocessed consistent with the PS cell libraries. Filtered counts matrices from the two 10x v3 chemistries (pbmc5k and NextGem) were downloaded from the online resource as they were already aligned to the same reference as the rest of the PS data. We note that the pairs of libraries from each technology were derived from the same biological donor (‘H1’ for v2 libraries; ‘H2’ for v3 libraries).

Utilizing the filtered gene by cell counts matrices for all scRNA-seq libraries, we identified and removed putative cell doublets using scrublet^74^ with the default parameters and specifying a 5% expected doublet rate. Barcodes identified as cell doublets were then filtered. Next, we performed data integration across these seven libraries for the PBMC reference projection via Azimuth via Seurat v4^28^. Differential gene expression summary statistics from scRNA-seq libraries were computed using edgeR^75^ while adjusting for sequencing technology (10x v2 or v3) and scaled number of genes detected per cell (see edgeRQLFDetRate^76^). We note that while edgeR was originally introduced for bulk RNA-seq, a comparison of differential expression tools demonstrated good performance for this approach compared to other bulk and single-cell strategies^76^. We performed gene-set enrichment analyses with the Panther Pathway enrichments using the WebGestalt framework^77^ using a rank ordering of genes by the signed Z score (**Fig. 3b**). Focused pathway enrichment analyses on metabolism pathways (**Extended Data Fig. 3f**) were performed using the clusterProfiler package^78^. Bulk expression data from GTEx was curated from the GTEx online portal for indicated genes^79^. All other visualizations and analyses for scRNA-seq data were performed using the Seurat framework.

### Chromosome 7 copy number (deletion) analysis

Chromosomal deletion 7q (del7q) analyses were performed only for PT3 as the other patients showed no evidence of copy number alterations (either from cytogenetics or sequencing data). To assign cells as either wild type or del7q, we performed copy number analyses of the accessible chromatin (scATAC-seq) data for all libraries and cells profiled from PT3. From the cytogenetics and sequencing data, we estimated the position 110,000,000 (on chromosome 7q22) as the approximate break point and computed the fraction of fragments occurring after this coordinate to get an estimate of the copy number changes across the various different libraries (shown in **Fig. 4b**). To call the single-cell del7q status, we used CONICS^36^ on the gene activity matrix and specified a custom region spanning the deletion for estimation of the copy number. Since the del7q abundance varied from between biological sources (e.g. PBMCs and CD34+ cells) and resulted in different maximum-likelihood estimates for the Gaussian distribution parameters, wild type or single-cell del7q genotype per-cell was called using a manual threshold of the predicted probability of the two-component mixture model based on the density of the first component predicted probability.

### Bone marrow ASAP-seq analyses

For the ASAP-seq antibody tag data, per-cell, per-antibody tag counts were enumerated via the kite | kallisto | bustools framework accounting for unique bridging events as previously described^25,80^. Cells called by the CellRanger-ATAC knee call were filtered based on abundance of protein (>150 unique molecules) and accessible chromatin (>1,000 nuclear fragments) as well as accessible chromatin enrichment (>25% fragments in accessibility peaks) and minimal non-specific antibody binding (<10 molecules associated with isotype control antibodies). Dimensionality reduction and clustering were performed only using the chromatin accessibility modality of the ASAP-seq and protein expression and gene activity values were used to annotated clusters as previously described^25^. Differential protein and gene activity score calculations (via Signac^19^) were performed using the FindMarkers function in Seurat. Somatic mtDNA mutations were identified by running mgatk on the ASAP-seq cells exceeding a mean 20x coverage using the default parameters^9^. For CD34+ analysis and projections (**Fig. 4d-g**), we utilized the LSI reference projection and reference CD34+ landscape as previously described^37^. All code needed to reproduce these analyses and embeddings is available as part of the online computational resource.

### *In vitro* erythroid culture single-cell analyses

Raw sequencing data was demultiplexed and aligned using CellRanger and CellRanger-ATAC as done for the PBMC analyses. Differential gene expression, via edgeR^75^, and pathway enrichment (Panther Pathway enrichments using the WebGestalt framework^77^) were conducted using the same workflow as the PBMC data. We computed a per-cell erythroid module score using 99 genes (e.g. *GATA1, ALAS2, HBB)* highly upregulated in erythropoiesis from our previous bulk transcriptomic atlas of cells derived from this *in vitro* system^81^ using the AddModuleScore function in Seurat.

## CODE AVAILABILITY

Software and documentation for mitochondrial DNA variant calling, including deletion calling and heteroplasmy estimation, is available via the mgatk package at http://github.com/caleblareau/mgatkas of version 0.6.1. All custom code to reproduce analyses is available at https://github.com/caleblareau/pearson_reproducibility.

## DATA AVAILABILITY

Data associated with this work is available at GEO accession **GSE173936** with reviewer access **tokencnotackcbbehbyx**.

